# k-Means NANI: an improved clustering algorithm for Molecular Dynamics simulations

**DOI:** 10.1101/2024.03.07.583975

**Authors:** Lexin Chen, Daniel R. Roe, Matthew Kochert, Carlos Simmerling, Ramón Alain Miranda-Quintana

**Affiliations:** Department of Chemistry, University of Florida, FL, USA; Quantum Theory Project, University of Florida, FL, USA; Laboratory of Computational Biology, National Heart, Lung, and Blood Institute, National Institutes of Health, Bethesda, Maryland, USA; Laufer Center for Physical & Quantitative Biology, Stony Brook University, Stony Brook 11794, USA; Department of Chemistry, Stony Brook University, Stony Brook 11794, USA; Department of Applied Mathematics and Statistics, Stony Brook University, Stony Brook 11794, USA

**Keywords:** algorithms, clustering, molecular dynamics, protein folding, *k*-means, conformational analysis

## Abstract

One of the key challenges of *k*-means clustering is the seed selection or the initial centroid estimation since the clustering result depends heavily on this choice. Alternatives such as *k*-means++ have mitigated this limitation by estimating the centroids using an empirical probability distribution. However, with high-dimensional and complex datasets such as those obtained from molecular simulation, *k*-means++ fails to partition the data in an optimal manner. Furthermore, stochastic elements in all flavors of *k*-means++ will lead to a lack of reproducibility. *K*-means *N*-Ary Natural Initiation (NANI) is presented as an alternative to tackle this challenge by using efficient *n*-ary comparisons to both identify high-density regions in the data and select a diverse set of initial conformations. Centroids generated from NANI are not only representative of the data and different from one another, helping *k*-means to partition the data accurately, but also deterministic, providing consistent cluster populations across replicates. From peptide and protein folding molecular simulations, NANI was able to create compact and well-separated clusters as well as accurately find the metastable states that agree with the literature. NANI can cluster diverse datasets and be used as a standalone tool or as part of our MDANCE clustering package.

## Introduction

Molecular Dynamics (MD) simulations serve as a computational microscope into intricate biological processes, yet the challenge arises when extending their scope to encompass longer timescales and larger systems. Unfortunately, the post-processing analysis of MD trajectories has struggled to keep pace with this demand, particularly evident in available clustering techniques, which is essential for unraveling protein dynamics and enhanced sampling techniques.^1^ Clustering, an unsupervised machine-learning method, identifies patterns within a dataset by organizing similar samples based on a similarity measure.^2,3^ When clustering structures obtained from MD simulations there are several metrics that can be used, but it is quite common to use root-mean-square deviation (RMSD) as the clustering metric.^4–6^ The RMSD is typically used in pair-wise manner, where the RMSD of a conformation is calculated to every other conformation in the ensemble, leading to *O*(*N*^2^) scaling of the clustering calculation. There are also many clustering algorithms, but choosing one often involves a stark trade-off: conventional algorithms such as *k*-means^7,8^ prove efficient but fall short in identifying subtle metastable states, whereas more robust methods like density-based clustering (e.g. DBSCAN,^9,10^ density peak^11^) incur significant computational overhead.^2,12,13^ In this contribution, we present the first module of our Molecular Dynamics Analysis with *N*-ary Clustering Ensembles (MDANCE) software package based on *n*-ary similarity (that is, using the notion of comparing multiple objects at the same time).^14–20^ Overall, MDANCE is aimed to provide great flexibility, new clustering algorithms, and novel tools to process the clustering results and to assess the overall quality of the clustering process. In the first installment of MDANCE, we will be introducing *k*-means *N*-ary Natural Initiation (NANI), an initialization method for *k*-means.

*K*-means is widely known to be the most popular clustering algorithm in the community for its straightforward approach. It was first proposed by Stuart Lloyd in 1957 and formalized by James MacQueen in 1967; now, it has become a central component of unsupervised learning for pattern recognition and data science. The input for *k*-means requires two parameters: the number of clusters and an initial estimate of the centroids. Beginning with initial centroid estimates, the algorithm proceeds by assigning each data point to the nearest centroid and recalculating the centroids for each cluster iteratively until convergence, in which the computed centroids remain the same, and data points stay within their respective clusters. When applied to large datasets from Molecular Dynamics, it proves to be efficient as it has a time complexity of *O*(*k* × *N* × *i*), *k* for the number of clusters, *n* for the number of points, and *i* for the number of iterations.^21^

Despite its efficiency, the *K*-means algorithm can struggle to identify metastable states due to one or more of the following issues: poor selection of the number of clusters (*k*), problems with the initial centroid estimation, and the clustering of non-convex shapes. The selection of *k* is a required input parameter for the algorithm. However, determining *k* can be challenging given the multidimensional and complex nature of most datasets, so the number of metastable states is often determined *a priori*.^22^ A larger *k* can result in finer partitioning of the data, which can identify some of the metastable states; however, this risks the accuracy of the cluster population and transition probabilities between clusters.^23^ The initial centroid estimation is also important because subsequent points are assigned based on these estimates. Therefore, many different initialization methods (e.g. *k*-means++^24,25^) were proposed to predict the best centroids for initialization (See Theory Section). However, all the existing centroid estimation algorithms are stochastic, which makes it challenging to assess the validity of the results. Lastly, *k*-means is not designed to identify non-convex cluster shapes since it is a partitioning cluster algorithm; the resulting cluster shapes tend to be uniform and globular and will usually not adapt arbitrary cluster shape.^26–29^ Another algorithm similar to *k*-means is *k*-medoids,^30^ which is similar to *K*-means except that the data points are assigned to the nearest medoid. Therefore, pairwise comparisons are performed to determine the similarity between every point to determine the least dissimilar object, making the algorithm less sensitive to outliers.^2^ Due to the pairwise comparisons and inefficiency with high-dimensional data, it has a time complexity of *O*(*k* × *N*^2^ × *i*).^31^ Since it is not as efficient as *k*-means, *k*-medoids will not be investigated in this study.

*N*-Ary Natural Initiation (NANI) is our proposed initialization technique for *k*-means with advantages of being fully deterministic, selecting the most diverse structures, and including a pipeline to determine the most optimal number of clusters. The main advantages of NANI are its efficient scaling, deterministic character, and ability to identify more compact and well-defined clusters over realistic MD simulations. Because all the initialization methods available are stochastic, NANI is an attractive option for producing reproducible results. A critical component of the NANI algorithm is diversity selection, which will be elaborated in the Theory Section. The extended similarity-based diversity selection can select a more diverse set of structures from a trajectory than *k*-means and hierarchical agglomerative clustering in *O*(*N*) time complexity.^18^ Lastly, the choice of number of clusters is not trivial, especially in a multidimensional dataset, where the user might even need knowledge about the system to make this decision.^32^ A pipeline has been established to make this decision to require minimal presumptions. A process involves screening through a range of *k* ‘s, and utilizing clustering quality metrics to determine the most optimal *k*. Through these additions, *k*-means clustering can obtain robust results even for multidimensional and complex datasets.

## Theory

### Tools

#### *N*-ary Similarity

Traditionally, we quantify the similarity of objects using pairwise functions, that is, taking two objects at a time. However, we have recently shown how it is possible to define *n*-ary functions to quantify similarity and differences,^14,15,17^ and that can take an arbitrary number of inputs at the same time. While most of the applications of these metrics have been in cheminformatic and drug design problems, here we use some of the basic principles to study conformational ensembles of different biomolecules.

#### Mean Squared Deviation

For a system with *M* atoms and *D* coordinates, the RMSD between the *i* th and *j* th frames, 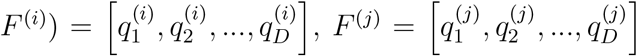 (where 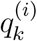 represents the *k* th coordinate of the *i* th frame), is calculated as:

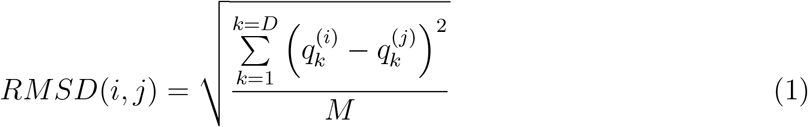

Similarly, we can define the Mean Squared Deviation (MSD) between two frames as:

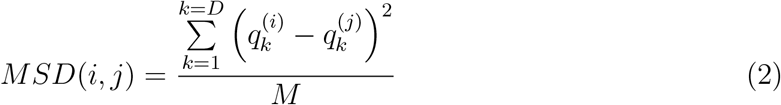

The key difference between these two magnitudes comes when calculating their average over *N* > 2 frames. For instance, in the case of the RMSD, it is clear that this will demand *O*(*N*^2^) scaling:

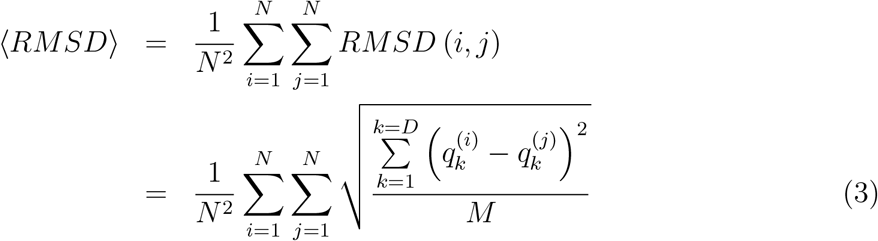

However, this is not the case for the MSD:

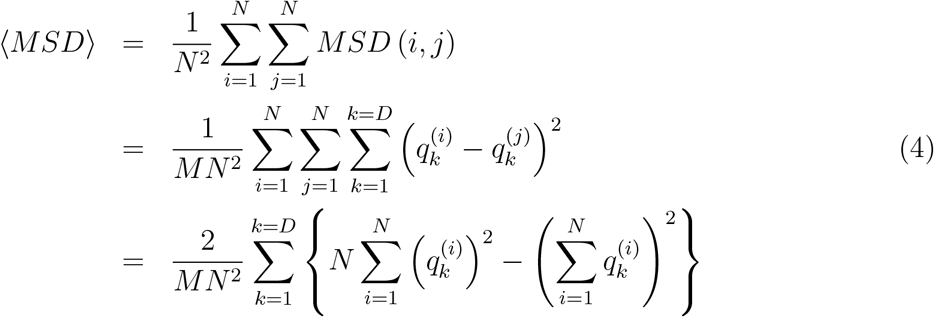

Notice that in the last equation, we only need 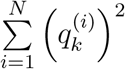, and 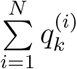 since they both scale as *O*(*N*), the overall MSD calculation has a much more attractive linear cost (while also bypassing the cumbersome square root calculation).

#### Complementary Similarity

Calculating the medoid of a set scales in *O*(*N*^2^), or *O*(*N* log *N*) at best. However, using complementary similarity, we can identify the medoid in *O*(*N*).^20^ The complementary similarity is based on the idea of the *n*-ary similarity; it calculates the *n*-ary similarity of a set without a given element. Therefore, the medoid will correspond to the lowest extended similarity as the set is the most dissimilar without the most representative member and the outlier will have the highest extended similarity. The complementary similarity will determine the ranking of how representative or outlier a member of the set is, from most representative (medoid) to most outlier-like.

#### Diversity Selection

Another useful tool in *n*-ary similarity is sampling representative structures from a set. It picks the first points using complementary similarity to find the medoid. It then picks the most dissimilar point from the selected point(s) at every iteration. That is, at any given point, we pick the point that minimizes the overall similarity of the selected set, as measured by an *n*-ary function.

#### Alignment

The most intuitive representation of an MD trajectory is the spatial coordinates of atoms in the trajectory. However, this can be challenging because configurations need to be aligned to some reference(s) to compensate for the effect of rotations and translations during a simulation and the decision of what reference(s) to align to can impact the data analysis significantly.^33,34^ Here, we explore two possibilities to handle the alignment: the traditional alignment to a single reference conformation, and the Kronecker version of the alignment proposed by McCullagh et al.^35^ Kronecker performs a global alignment and distributes weight according to the variance of the atoms’ coordinates, in essence, giving more weight to the more rigid areas of a biomolecule.

### Clustering

*K*-means is a partitioning clustering algorithm that assigns points to the closest centroid. It iterates by recalculating the centroid and reassigning points. Other than the determination of the number of clusters, initial centroid prediction is critical as the clustering is susceptible to modifications in the labeling with different centroid estimations. Therefore, a diverse set of initial centroids is required for better results than random initialization. Some of the available initialization methods, including our proposed NANI, are presented below:.

**Vanilla *k*-means++** is the original version of *k*-means++.^24^ The algorithm begins by initially picking a random point from the set. It will then pick the next point using a weighted probability distribution of distance from the selected point and will favor selecting the point with the maximum distance from the selected point. It will continue for *k* number of clusters. Since the algorithm has stochastic elements, results will vary every time.

**Greedy *k*-means++** follows the same algorithm as its vanilla version, except it runs multiple local trials of initialization and picks the best one for *k*-means clustering. The greedy version can perform better than the vanilla variant, but this is not guaranteed to be always the case^24,36^ and it is the default setting for initializing *k*-means in scikit-learn.^37^ This algorithm provides good results for simple and well-separated systems. However, its limitation is apparent in noisier data (See Fig. 2).

**Figure 1:**
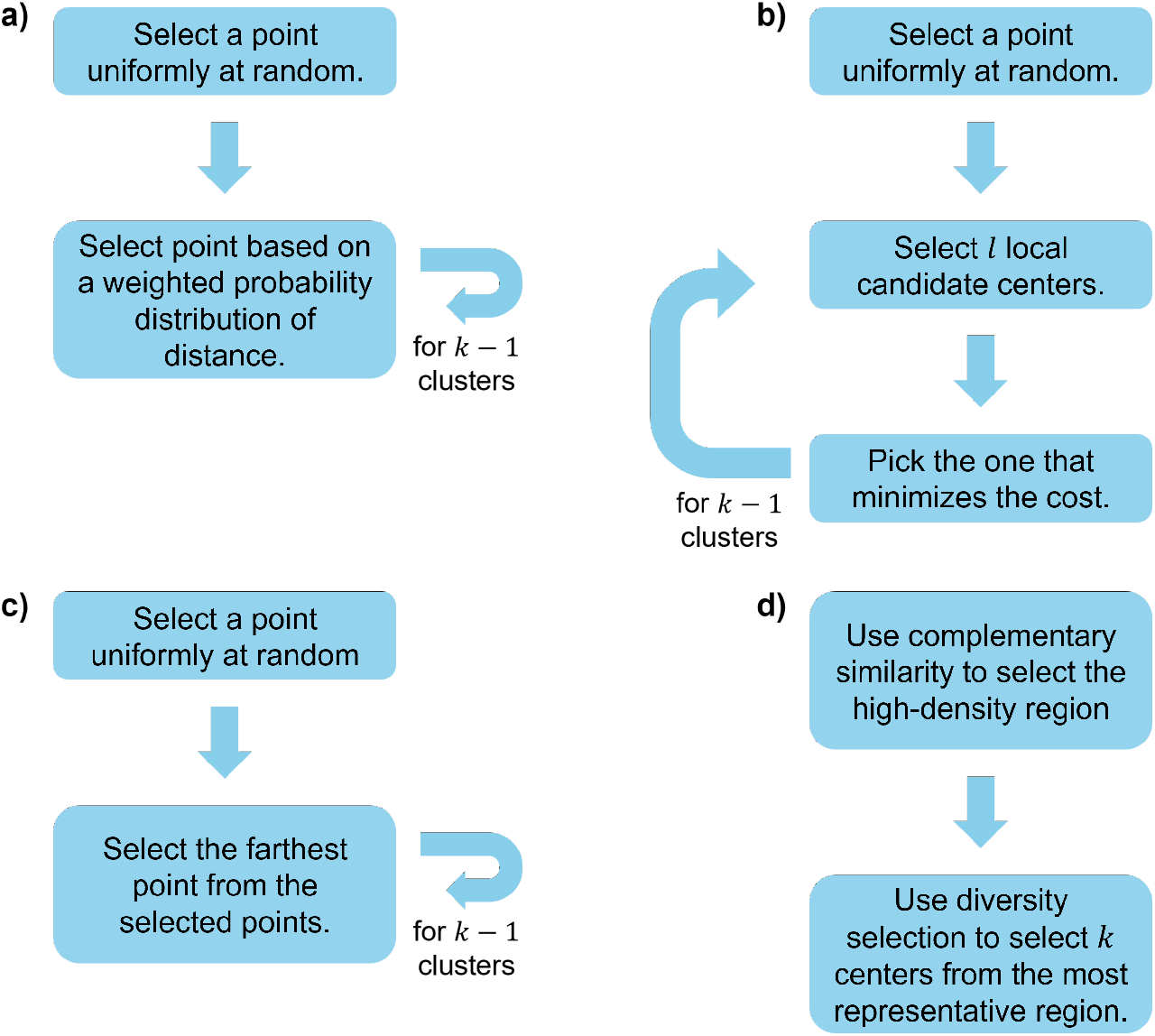
Four centroid selection algorithms for *k*-means initialization. **(a)** Vanilla *k*-means++ **(b)** Greedy *k*-means++ **(c)** CPPTRAJ **(d)** NANI

**Figure 2:**
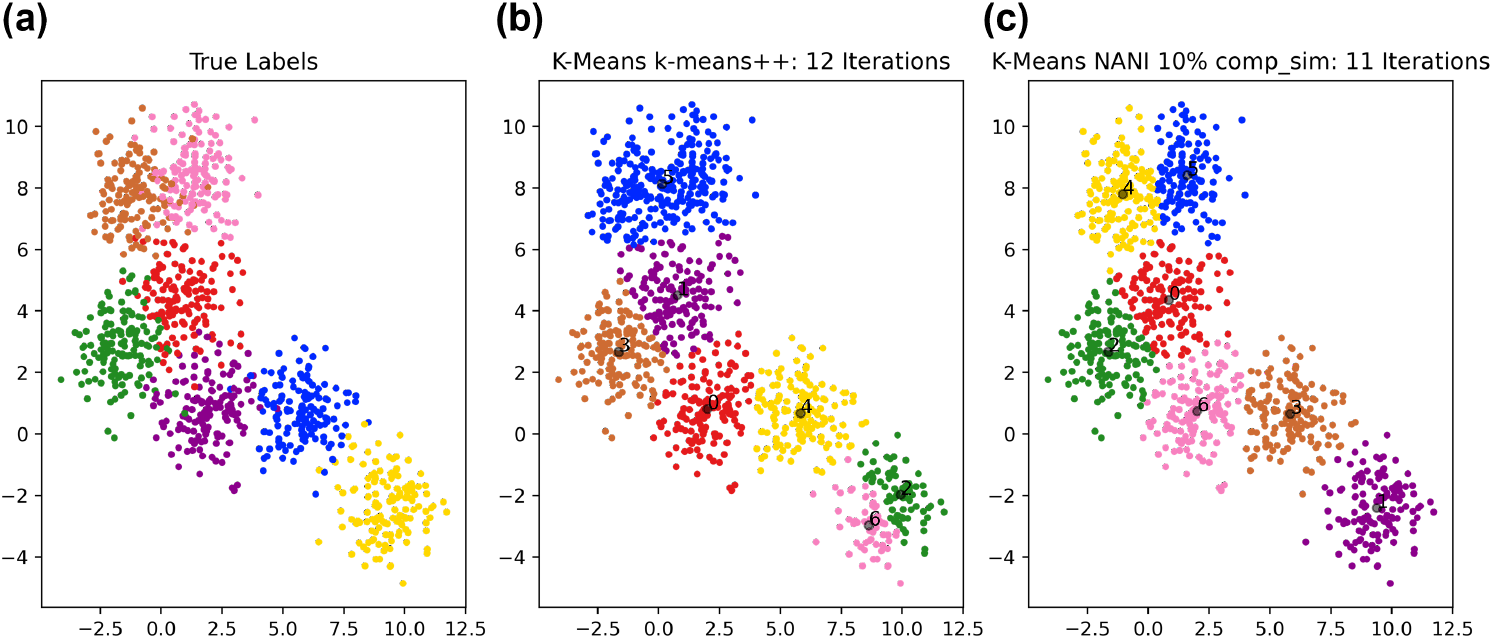
*k*-means NANI on a sample 2D data. A different color represents a different cluster label. The black dot represents the centroid of that cluster. **(a)** True Labels. **(b)** *k*-means clustering with centroids initialized by *k*-means++. **(c)** *k*-means clustering with centroids initialized by *k*-means NANI.

**CPPTRAJ Initialization** starts from a random point and selects the farthest point from the selected points.^38^ It then iterates for *k* number of clusters.

***N*-Ary Natural Initiation (NANI)** NANI first uses the medoid algorithm to stratify the data into high-density regions and then performs a diversity selection (`Max nDis`, `ECS MeDiv`) to pick well-separated centroids.^15^ The diversity selection starts with the medoid of the dataset. It iterates the data to find the object with the lowest similarity between the candidate and the existing set of centroids. This procedure is inspired by *k*-means++, but with the key difference that the first point is chosen in a completely deterministic way. MDANCE includes two NANI variants: div select is the closest one to the standard *k*-means++, in the sense that the last *k* − 1 centroids are chosen from all the remaining points in the set. On the other hand, the `comp sim` version of NANI is even more efficient, since we use all the complementary similarity values to stratify the data, and then do the diversity selection of the remaining *k* − 1 centers only over a high-density fraction of the initial points. A key insight here is that using the *n*-ary complementary similarity algorithm to stratify the data ensures that this algorithm retains the attractive scaling of traditional *k*-means.

## Systems and Software

### 2D Datasets

The model 2D data was were obtained from scikit-learn single-labeled generated dataset. The blob disk, nine diamonds, and ellipses were extracted from Veenman *et al*, Salvador *et al*, and Bandyopadhyay *et al*, respectively.^39–43^

### *β*-Heptapeptide

The topology and trajectory files correspond to publicly available data and were assessed through GitHub.^44,45^ The atom selection follows Daura *et al*, ^46^ Lys2 to Asp11, with N, C*α*, C, O, and H atoms. The terminal and side chain residues were ignored to minimize noise in the clustering. The single-reference alignment was done after aligning to the 1000^th^ frame. Also, as a comparison, we performed the Kronecker alignment.

### *β*-Hairpin

This system was adapted from a stabilized *β*-hairpin system proposed here.^47^ Orn-8 was substituted with Lys-8 due to lack of ornithine parameters in the force field; this substitution was tested previously^48^and was found to have minimal impact on the conformational stability. Subsequent MD simulations indicated that the peptide was often kinetically trapped due to strong salt bridging interactions between Arg-1 and Glu-4; for the purpose of facilitating ensemble convergence, we introduced an R1Q substitution. Neutral capping groups (ACE, NHE) also were introduced at the termini to further reduce the possibility of strong electrostatic traps. This modified system was then relaxed to stabilize the initial structure before eight independent MD simulations of ∼9 μs each were generated at 420K (∼70 μs total). The ensuing ensemble was very diverse, with backbone RMSD’s ranging from under 1 angstrom to over 10 angstrom and featuring ∼11,000 folding and unfolding events. ∼65,000 equally spaced coordinate sets were extracted for analysis. All clustering on this system used the backbone atoms (C*α*, C, and N) and all frames were aligned to the 1000^th^ frame that was extracted.

### HP35

Analysis was performed on a 305 *μ*s, all-atom simulation of Nle/Nle mutant of the C-terminal subdomain of the Villin headpiece (also known as HP35) from D. E. Shaw Research.^49^ This simulation was conducted at a temperature of 360 K and consists of 1.52 million frames with a frame separation of 200 ps. All frames in the simulation were aligned to the 5000^th^ frame in the simulations and frames before the 5000^th^ frame were discarded as a relaxation phase. The backbone atom selection encompassed the following atoms: N of residue 1, CA, C, N of residues 2 to 34, and N from residue 35; this selection is consistent with previous studies of HP35. A sieve was applied on every 20^th^ frame, resulting in a total of ∼89,000 frames for clustering.

### NuG2

An analysis was conducted on a dataset comprising four independent all-atom simulations of NuG2, a mutant variant of Protein G. The simulations, with a cumulative duration of 1.15 milliseconds (5.78 million frames with a frame separation of 200 ps), were conducted at a temperature of 350 K and carried out by D. E. Shaw Research.^50^ All frames in the simulation were aligned to the 5000^th^ frame in the simulations and frames before the 5000^th^ frame were discarded as a relaxation phase. The atom selection encompasses the backbone atoms (CA, C, and N) of residues 1 to 56; this selection is consistent with previous studies of NuG2. A sieve was applied on every 65^th^ frame, resulting in a total of ∼76,000 frames for clustering.

### MDANCE

The NANI code is available as one of two modules in the MDANCE package. MDANCE contains basic functionality in a Behind The Scenes (BTS) module that handles I/O functions, interfacing with packages like AMBER^51^ and MDAnalysis,^52,53^ along with several alignment options. The NANI module handles the initial seed selection procedure for both the `comp sim` and `div select` variants, but it can easily be modified to include other selection criteria. While the applications in this paper are centered around the use of the MSD as *n*-ary difference measure, MDANCE gives the user the option to use from other *n*-ary indices, like extended versions of popular cheminformatic metrics like Tanimoto, Russel-Rao, etc. The MDANCE GitHub repository can be found here: https://github.com/mqcomplab/MDANCE.

## Results

### 2D Datasets

Clustering evaluation metrics assess the quality of the resulting clusters. Clustering evaluation metrics are separated into internal and external measures; internal measures do not use ground-truth labels, whereas external measures require ground-truth labels. Internal measures are primarily used for complex data where a ground-truth label is unavailable. Several indicators were studied to identify the most optimum metric for identifying the number of clusters in a dataset.

V-measure score^54^ calculates how accurately the clustering labels match the ground truth based on two criteria—homogeneity and completeness. The dataset is divided into classes and after clustering, the algorithm separates the dataset into clusters. Homogeneity measures the purity of the cluster if all the members in a cluster contain only members with the same class label. Completeness measures if all the members with the same class label are in one cluster. These two criteria will calculate a V-measure score between 0 to 1. A V-measure closer to 1 indicates greater accuracy to the ground truth labels. This metric is an external measure, which requires ground truth labels as input. Therefore, the V-measure will not be used for simulation data since there are no known cluster labels.

Calinski-Harabasz index^55^ (CH, also known as variance ratio criterion) is a clustering evaluation metric that quantifies the separation between clusters and the compactness within clusters. CH is a ratio of between-cluster dispersion and within-cluster dispersion. Between-cluster dispersion is a sum of squares between each cluster centroid and the centroid of the dataset. Within-cluster dispersion is the sum of squares between samples in the data and their respective cluster centroids. A high CH value indicates well-separated clusters. CH index is an internal measure, which is solely based on the dataset and clustering results and does not use ground-truth labels in the calculation, which makes it a candidate for a metric for quantifying simulation data.

Both the Calinski-Harabasz index (CH) and Silhouette score are popular cluster evaluation metrics for measuring cluster quality in the community.^1,32^ Silhouette score^56^ is also an internal clustering measure and calculates intra- and inter-cluster distances. For every data point, the algorithm calculates the average distance to all other points in that cluster (*a*) and also calculates the average distance to all the points in the nearest cluster (*b*). The score is calculates using *silhouette score* = (*b* − *a*)*/* max(*a, b*). Values range between -1 to 1, with -1 meaning the point is clustered poorly, values close to 0 meaning the point is close to a border with another cluster, and 1 meaning the point is well-clustered. Because silhouette score typically requires a pairwise matrix, it is *O*(*N*^2^) and the slowest out of all mentioned metrics.

Average mean squared deviation (MSD) only takes into account the clustering labels and the coordinates and determines how compact clusters are, in which a lower MSD would be a tighter cluster. Similar to RMSD, mean squared deviation determines the similarity between frames. Average MSD calculates the similarity of clusters using the *n*-ary framework. This metric is the least computationally intensive out of all the clustering quality metrics.

Similar to CH and Silhouette score, the Davies-Bouldin index (DBI)^57^ is an internal clustering measure and quantifies similarity within and between clusters. DBI is a ratio between intra-cluster and inter-cluster similarity. The intra-cluster similarity is the average distance between points in a cluster and the centroid of the cluster. The inter-cluster similarity is the distance between the cluster centroids. A lower DBI value indicates clusters that are far apart and tightly packed.

Not only is *k*-means NANI (`comp sim` and `div select`) both a deterministic seed selector, but it also results in more compact and well-defined clusters with linear scaling comparable to *k*-means++. In our 2D results, *k*-means NANI was able to correctly identify the clusters of noisy data (See Fig. 2) because it begins with the medoid and only selects from high-density regions. Fig. 3 shows measures of the robustness of different clustering techniques using average MSD, CH index, DBI index, number of iterations, and V-measure score. `div select` is similar to the initialization used in CPPTRAJ because both choose the farthest point from the already selected points. Since `div select` only sometimes surpasses the other variants and usually takes more iterations compared to `comp sim`, only `comp sim` will be investigated in the simulation data. On the other hand, `comp sim` surpassed *k*-means++ variants in all clustering metrics, further motivating its use as a seed selector for *k*-means clustering. Therefore, from now on, whenever we refer to NANI, we are implicitly referring to the `comp sim` variant.

**Figure 3:**
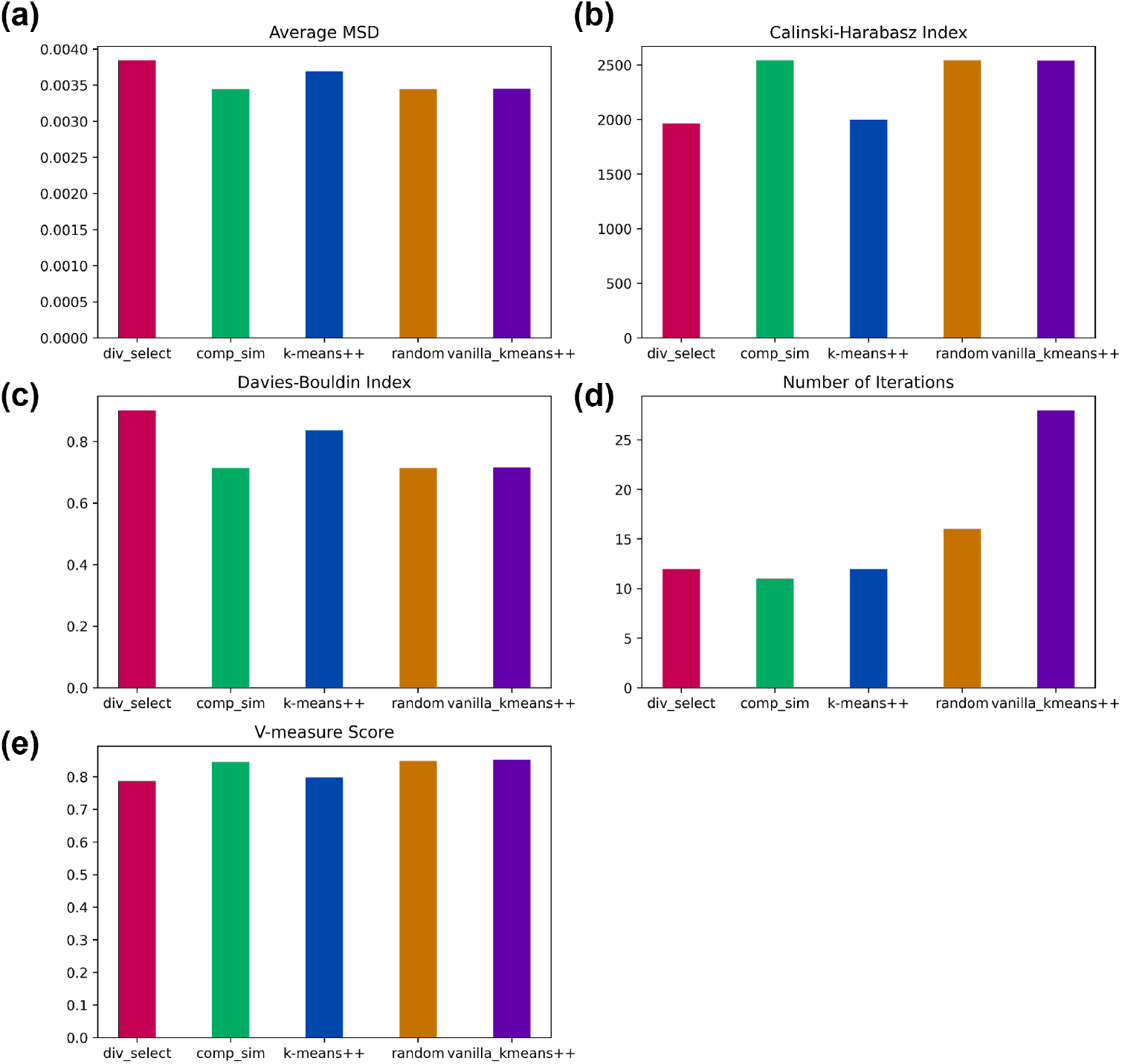
Cluster characteristics of different seed selectors on a sample 2D data from Fig. 2. **(a)** Average MSD of the clusters using different seed selectors. **(b)** Calinski-Harabasz index. **(c)** Davies-Bouldin index. **(d)** Number of iterations. **(e)** V-measure score.

### Application to Peptide Systems

A key part of our clustering pipeline is the incorporation of the previously discussed indicators to quantify the quality of the final clustering. This is critical to identify the optimum number of clusters in the data. In all cases, we performed a scan over different values of *k*, which is especially necessary when analyzing realistic MD simulations. Then, different clustering quality metrics will be explored to identify the most ideal metrics for the clustering pipeline hereinafter.

The Calinski-Harabasz index (CH) measures how well-separated clusters are and the trend for CH is that a larger index would indicate an optimal number of clusters. However, all the graphs exhibit the same behavior as shown in Fig. 4 with changing different variables (systems, alignment method, initialization technique). It seems unlikely that two clusters would be the most optimal number of clusters in every case; however, CH tends to favor a small number of clusters due to its bias for convex shapes. Overall, the behavior of NANI and the two *k*-means++ variants is very similar, with the CPPTRAJ algorithm consistently resulting in lower CH values. Also note the variations on the CPPTRAJ and *k*-means++ results, especially for the *β*-hairpin and HP35 systems. As far as the quality of the clusters goes, there does not seem to be an increase in the separation of clusters when we use the Kronecker alignment (with the *β*-heptapeptide and *β*-hairpin even showing a better behavior for the single reference alignment).

**Figure 4:**
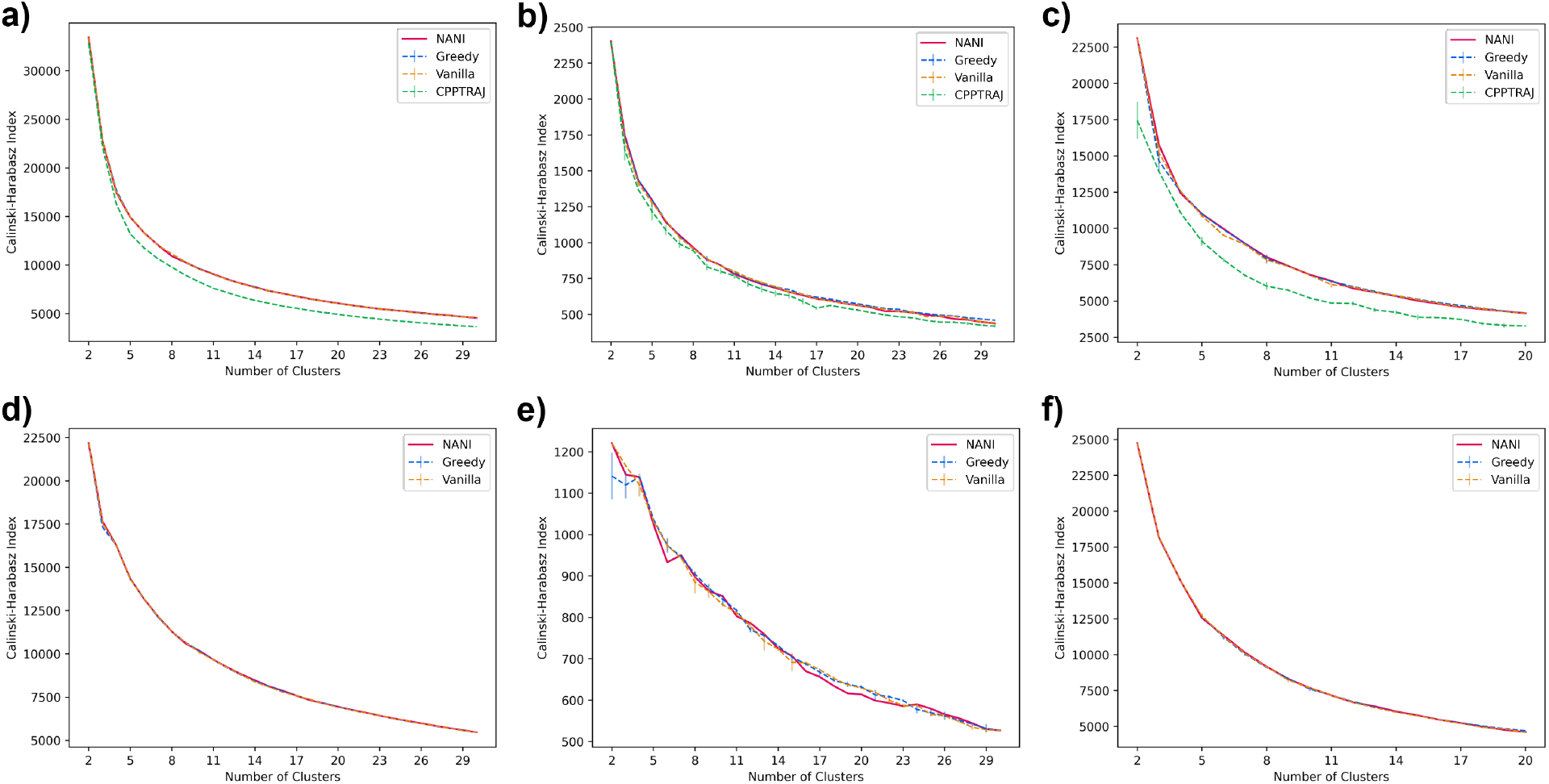
Calinski-Harabasz index (y-axis) vs Number of clusters (x-axis) of different seed selectors applied on three peptide systems. Error bars represent the standard deviation over three replicates. **(a)** *β*-hairpin single-reference aligned **(b)** *β*-heptapeptide single-reference aligned **(c)** HP35 single-reference aligned **(d)** *β*-hairpin Kronecker aligned **(e)** *β*-heptapeptide Kronecker aligned **(f)** HP35 Kronecker aligned

**Figure 5:**
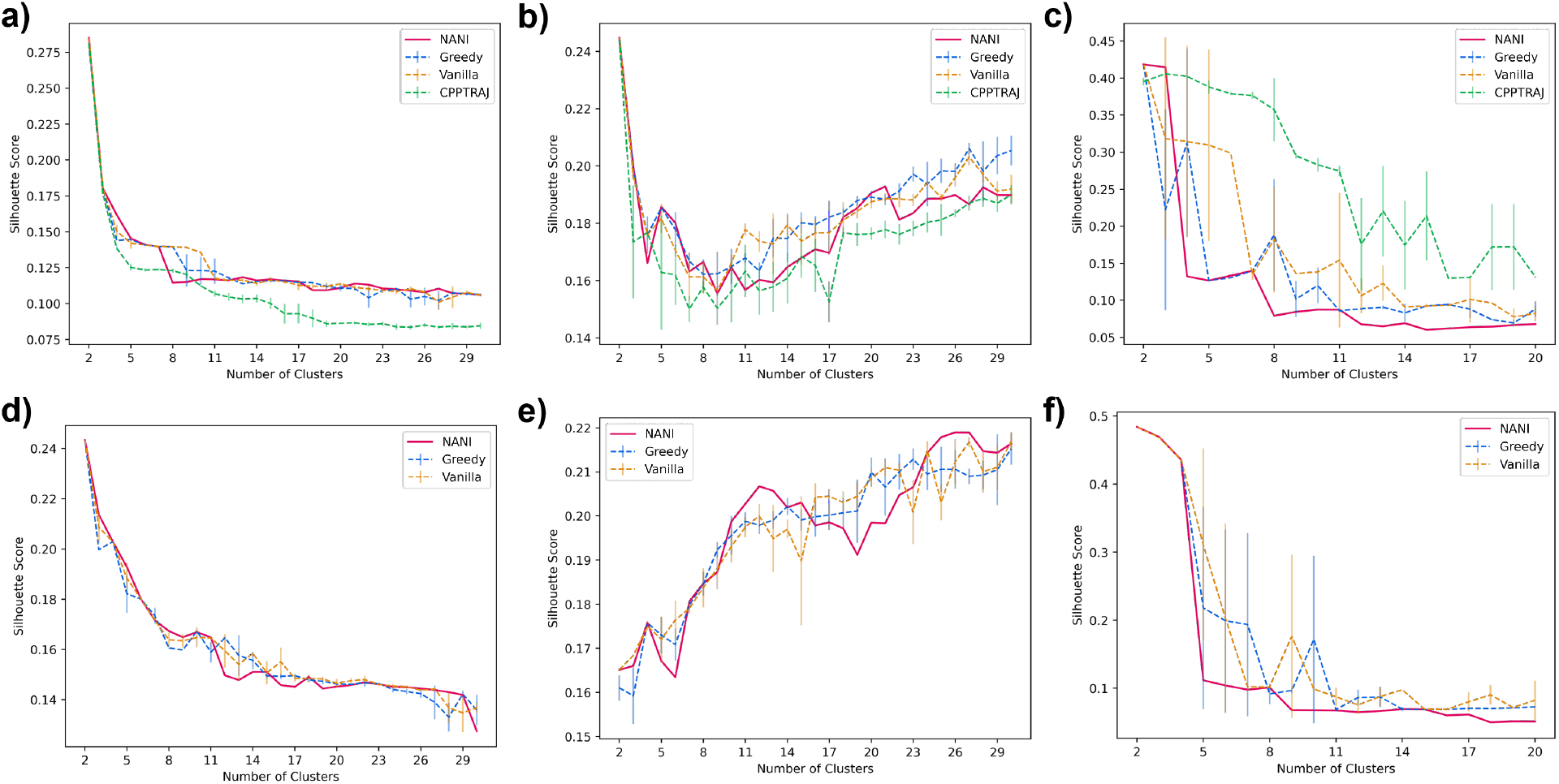
Average Silhouette score (y-axis) vs Number of clusters (x-axis) of different seed selectors applied on three peptide systems. Error bars represent the standard deviation over three replicates. **(a)** *β*-hairpin single-reference aligned **(b)** *β*-heptapeptide single-reference aligned **(c)** HP35 single-reference aligned **(d)** *β*-hairpin Kronecker aligned **(e)** *β*-heptapeptide Kronecker aligned **(f)** HP35 Kronecker aligned

**Figure 6:**
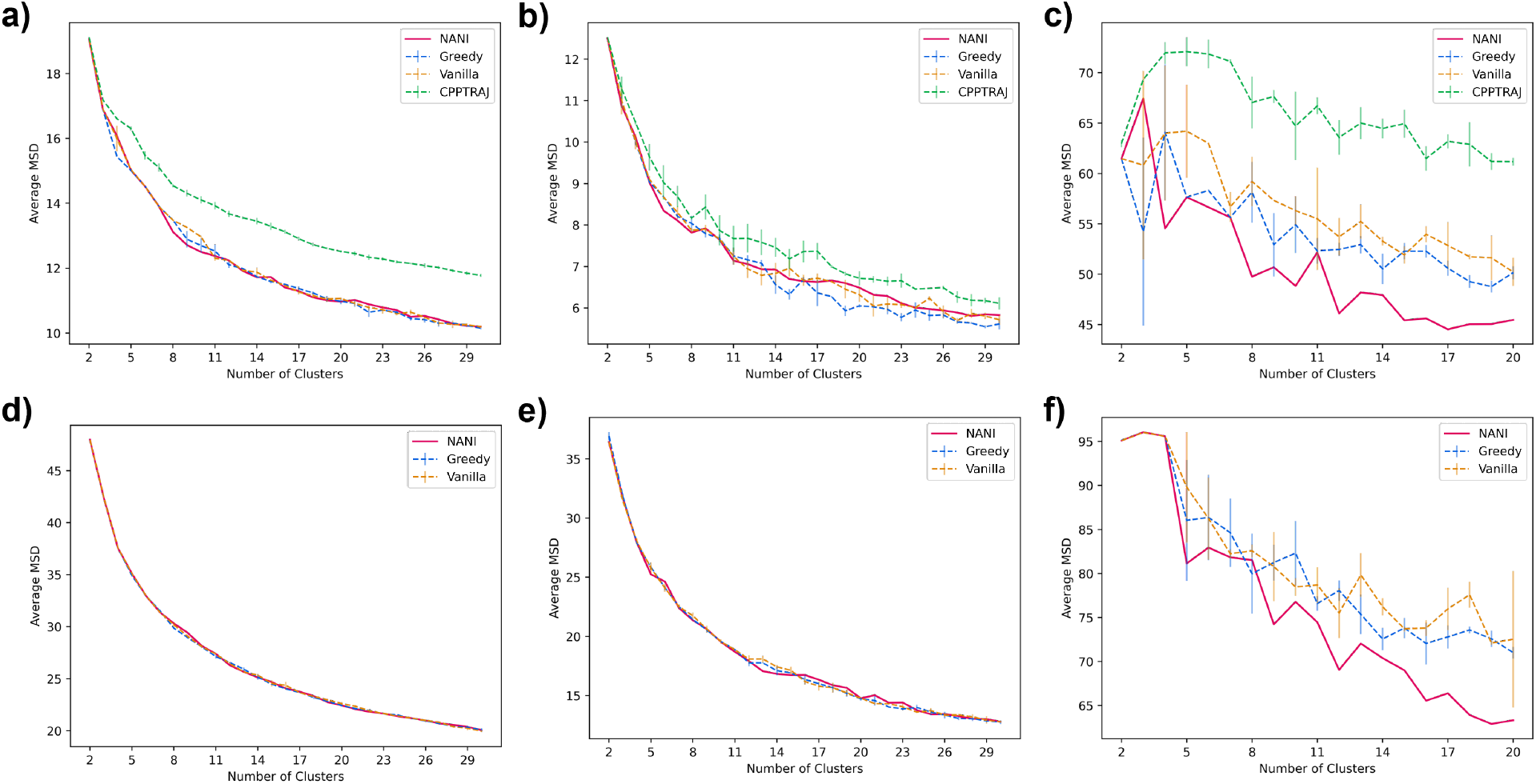
Average mean squared deviation (y-axis) vs Number of clusters (x-axis) of different seed selectors applied on three peptide systems. Error bars represent the standard deviation over three replicates. **(a)** *β*-hairpin single-reference aligned **(b)** *β*-heptapeptide single-reference aligned **(c)** HP35 single-reference aligned **(d)** *β*-hairpin Kronecker aligned **(e)** *β*-heptapeptide Kronecker aligned **(f)** HP35 Kronecker aligned

The Silhouette score is another metric to measure similarity within and between clusters. A similar issue with this metric is that it favors a smaller number of clusters due to its bias for convex clusters. A bigger issue with the Silhouette and CH indices is that their change from one *k* value to another does not seem to be a good indicator of the optimum number of clusters in the set. This is particularly evident in the case of CH, with the values just monotonically increasing with decreasing *k* values. A similar pattern is followed by the Silhouette score, which is more evident for the *β*-heptapeptide and HP35 cases. There, the score just tends to either remain essentially constant over a range of *k* values or just tends to increase. The *β*-hairpin system highlights a particularly pathological Silhouette behavior, with the single reference and Kronecker alignments presenting totally opposed trends for this score. Given that the *β*-hairpin simulations were conducted at the highest temperatures, and thus are expected to boast a higher conformational plasticity, this seems to indicate that the silhouette score is not appropriate to identify the optimum *k* values for these simulations.

The average mean squared deviation (MSD) would indicate how compact a cluster is. MSD follows the trend of a decreasing concave-up shape due to the nature of an increasingly more dissimilar cluster as it grows and a more similar cluster as it is smaller. Just like for the CH index, the MSD values tend to change monotonically, so at first sight, they do not seem suitable to identify optimum *k* values. However, interestingly, in several cases, we could correlate the changes in the slope of the MSD graphs with the behavior of optimum *k* values of other indicators, chiefly, the Davies-Bouldin index analyzed in the next section. When NANI was applied to the Molecular Dynamics trajectory, the compactness was greater than that of *k*-means++ as demonstrated by the average MSD values.

Davies-Bouldin index (DBI) is the metric that works the best for the most number of systems. DBI measures both the intercluster and intracluster distance and a lower DBI would indicate an optimal number of clusters. This is usually the case for simple systems, but with high-dimensional and complex datasets this can sometimes be misleading because DBI tends to favor a small number of clusters, the minimum value is not always the optimum number. Moreover, for most indicators designed to quantify the quality of the clustering, it can be argued that their local behavior is more important than their absolute values.

(This has been recently emphasized by Hocky, McCullagh, *et al*.^58^) While this is in principle true for any of these indices (and, in a sense, can be seen as the rationalization behind the “elbow method”), it is more useful in the case of the DBI. Once again, the almost perfectly monotonic behavior of the CH index renders this approach useless. On the other hand, the quasi-constant behavior of the Silhouette index over intervals of *k* values, coupled with an almost monotonic increase for some systems, also makes this a relatively difficult criterion to adapt in that case. Therefore, we introduced another criterion for measuring the optimal DBI—the maximum second derivative DBI. The maximum 2^nd^ derivative of DBI with respect to the number of clusters is the local minimum of the two steepest slopes, which can often indicate the most optimal value before it rises and drops back to global minima, which is due to DBI’s bias for convex clusters. In other words, the 2^nd^ derivative criterion allows users to distinguish “unusually stable” *k* values, while tending to be more robust towards the bias for particular cluster shapes. From Fig. 7, DBI proves to be a more robust metric compared to CH and silhouette score as the topology of the graph is different for different systems, while also remaining easily quantifiable, and without presenting obvious biases towards a particular range of *k* values. Comparing the DBI for the different initialization methods, the CPPTRAJ method gives the highest DBI would indicate a weaker initialization method than the other three since it is farthest from the desired low values. However, both vanilla and greedy *k*-means++ methods were not as robust as NANI as there were great fluctuations in the DBI values between runs. Since all methods contain elements of probability and randomness, they are not as robust as the deterministic NANI methods because they will give a different optimal number of clusters every time the user executes the methods. This pathological behavior is shown in Table 1, where we present the results of three replicates of Greedy, Vanilla, and CPPTRAJ *k*-means. Notice that for all the peptide systems, and independently of if the optimum *k* is selected with the absolute value or the 2^nd^ derivative of DBI, different runs tend to give different values for the optimum value of *k*. NANI not only gives the DBI values in every replicate, but it also gives DBI values of the same quality, if not better in several instances, than those of the *k*-means++ variants.

**Table 1:**
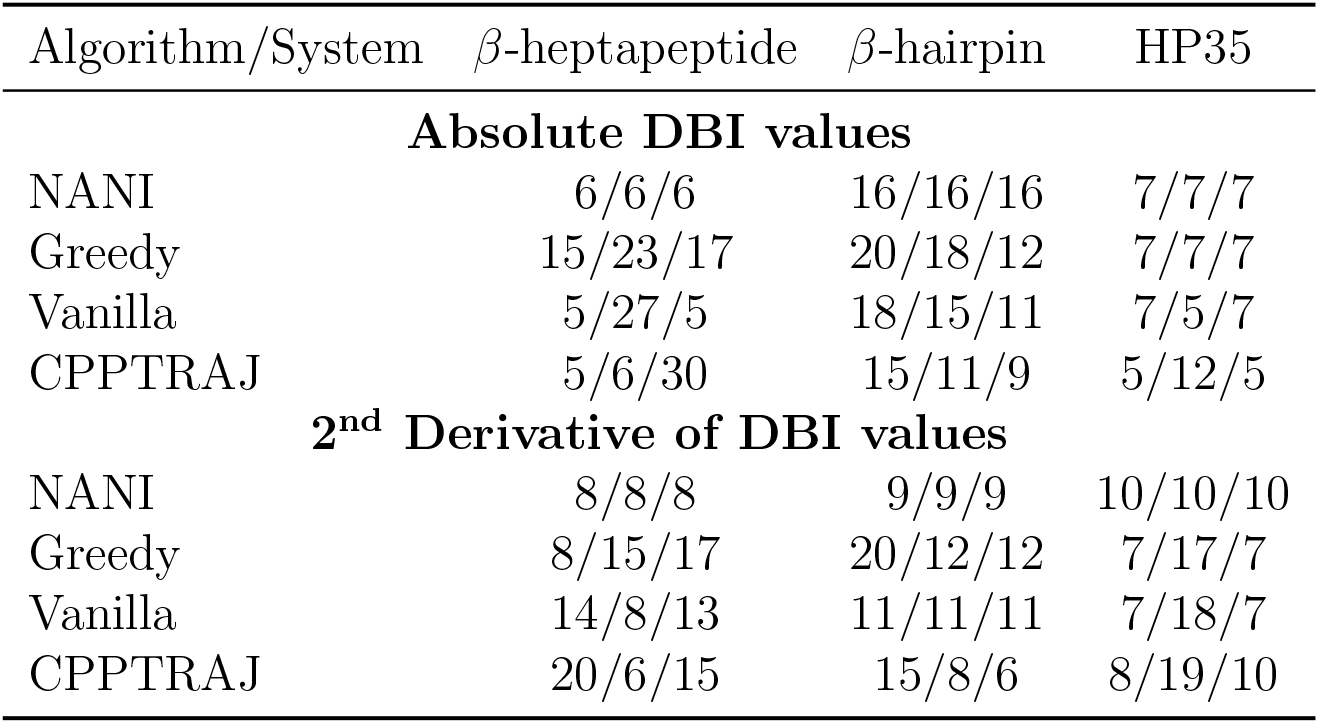
Optimum number of clusters over for the studied systems, as determined by running three replicas of the Greedy, Vanilla, and CPPTRAJ algorithms. Top: *k* values determined from the absolute minimum of the DBI index. Bottom: *k* values determined from the 2^nd^ derivative of the DBI index. In all cases, only *k* values bigger or equal to 5 were considered.

**Figure 7:**
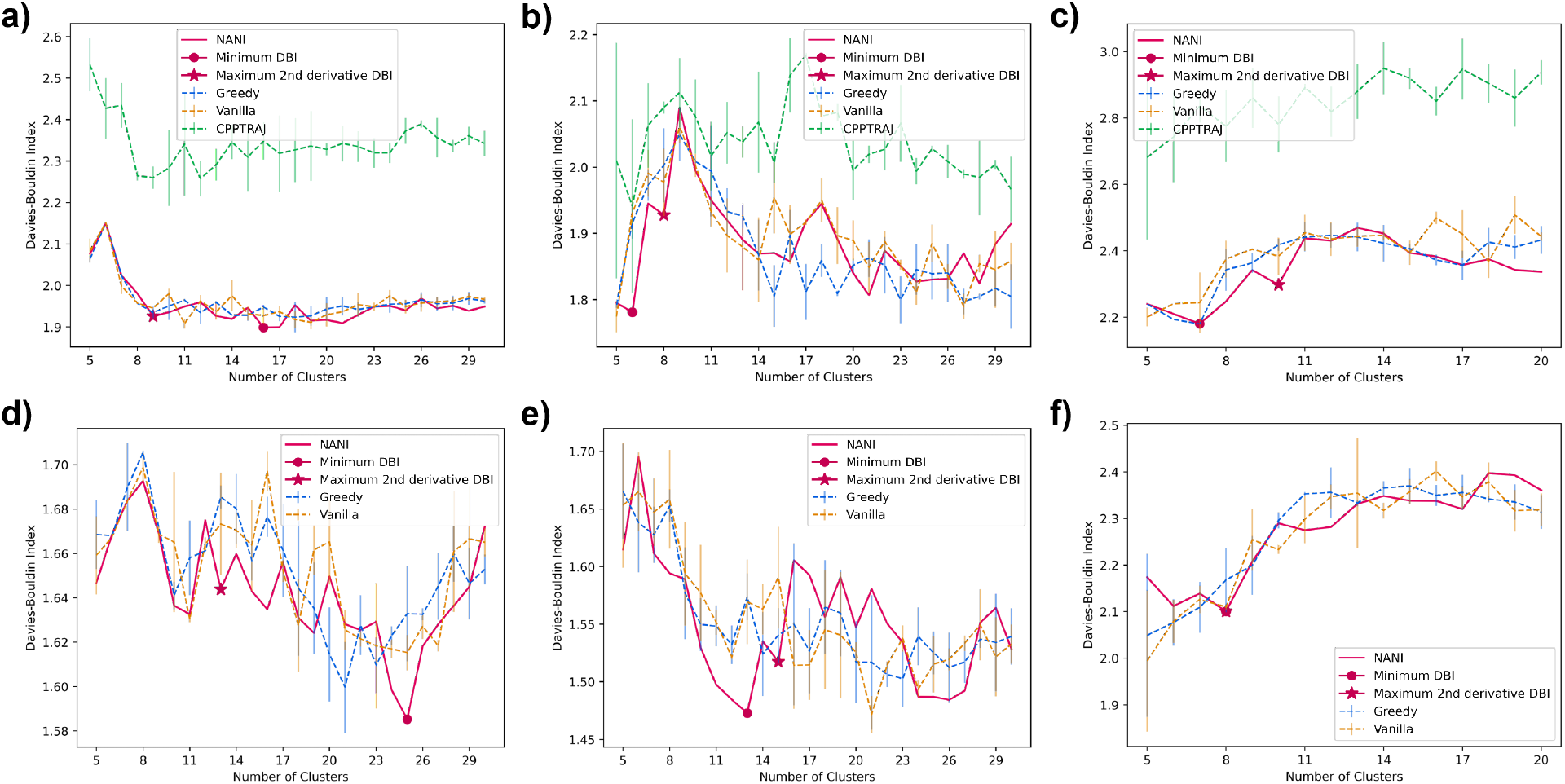
Davies–Bouldin index (y-axis) vs Number of clusters (x-axis) of different seed selectors applied on three peptide systems. Error bars represent the standard deviation over three replicates. **(a)** *β*-hairpin single-reference aligned **(b)** *β*-heptapeptide single-reference aligned **(c)** HP35 single-reference aligned **(d)** *β*-hairpin Kronecker aligned **(e)** *β*-heptapeptide Kronecker aligned **(f)** HP35 Kronecker aligned

To further highlight NANI’s performance, we dissected the HP35 simulations in more detail, given the recent interest in this system as a benchmark for other clustering methods. In the application to HP35, the number of clusters was determined to be seven using single-reference alignment and calculated using NANI as the seed selector. In the overlaps (See Fig. 8, it shows that NANI was able to identify seven metastable states of HP35 with cluster populations consistent with a previous study (using more computationally demanding clustering methods) in which six metastable states were reduced to a four-state model consisting of a native state N (53% of the population), a native-like state N’ (14%), an intermediate state I (18%), and an unfolded state U (15%).^35^ NANI obtains similar populations for N (66%), N’ (8%), I (9%), and U (17%). The results are consistent for both the “condensed” version of the results (corresponding to *k* = 4), or the optimal configuration with *k* = 7. N’ is the partial unfolding of helix 3.^35,59^ In all cases, the N’ conformation was found to be the least populated. N1 and N2 are two conformations in the folded state (N) except for the orientation of helix 1, but both are highly tight clusters, representative of the native folded structure of HP35; overall, the N state is the most populated cluster. Three clusters corresponded to the unfolded state (U), with uncoiling observed in U1, U2, and U3. U2 has a unique orientation of helix 1 and 3 with both turning away from helix 2. U1 has a more defined structure in helix 3 than in U3. The intermediate structure is observed to have decreased variance from the unfolded states, with defined structures in all three helices but not yet completely folded.

**Figure 8:**
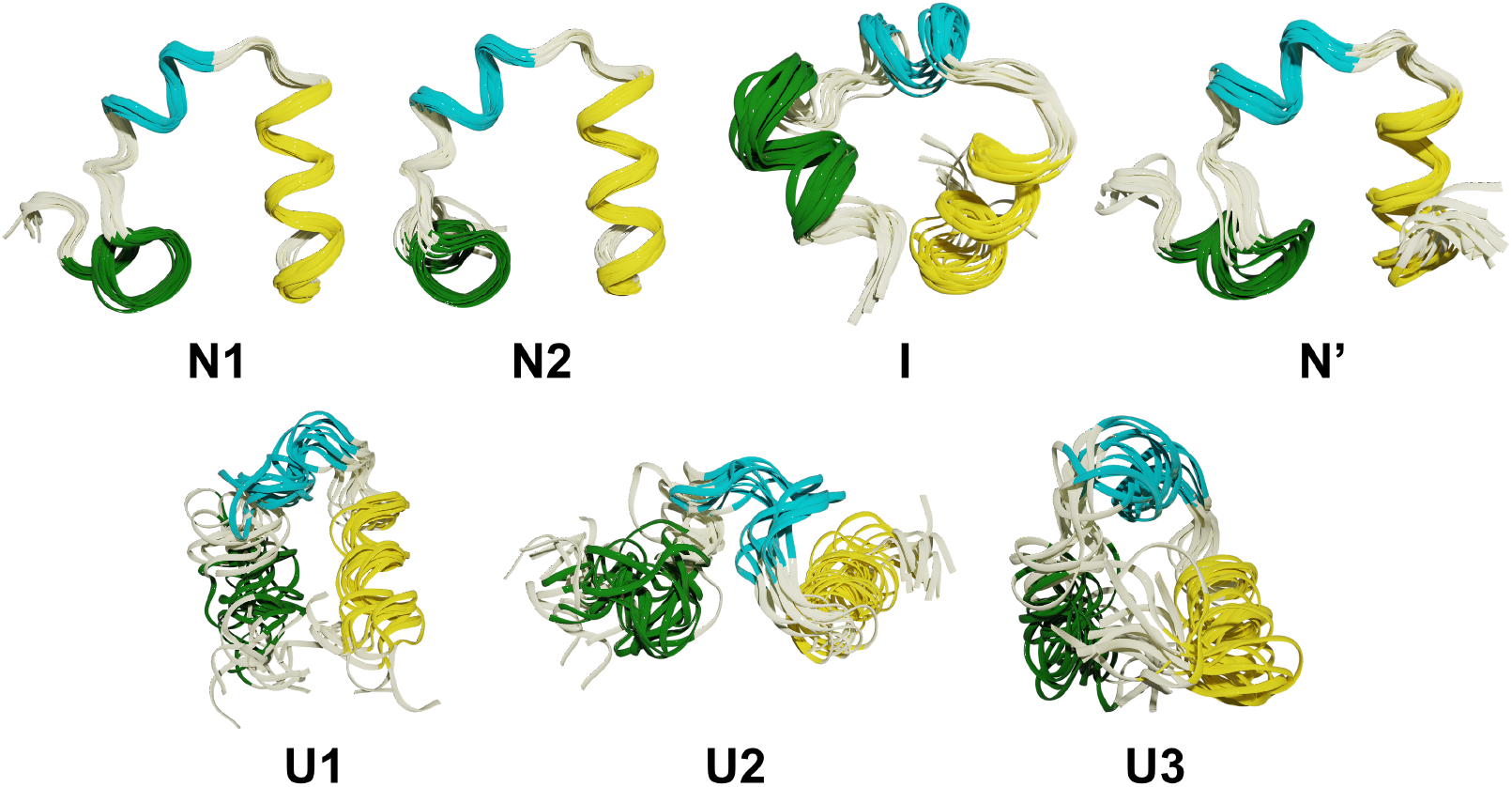
Structural overlaps for HP35 in four states: folded (N), partially folded (N’), intermediate (I), and unfolded (U). Helix 1, 2, and 3 are in green, cyan, and yellow, respectively.

**Figure 9:**
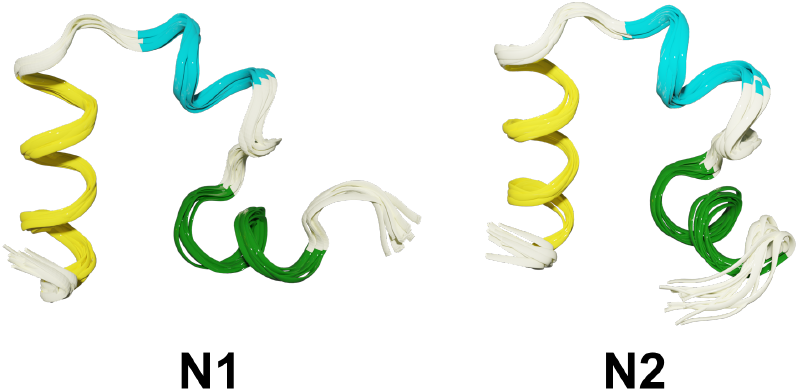
Another view of the structural overlaps for HP35 in two folded (N) states.

**Figure 10:**
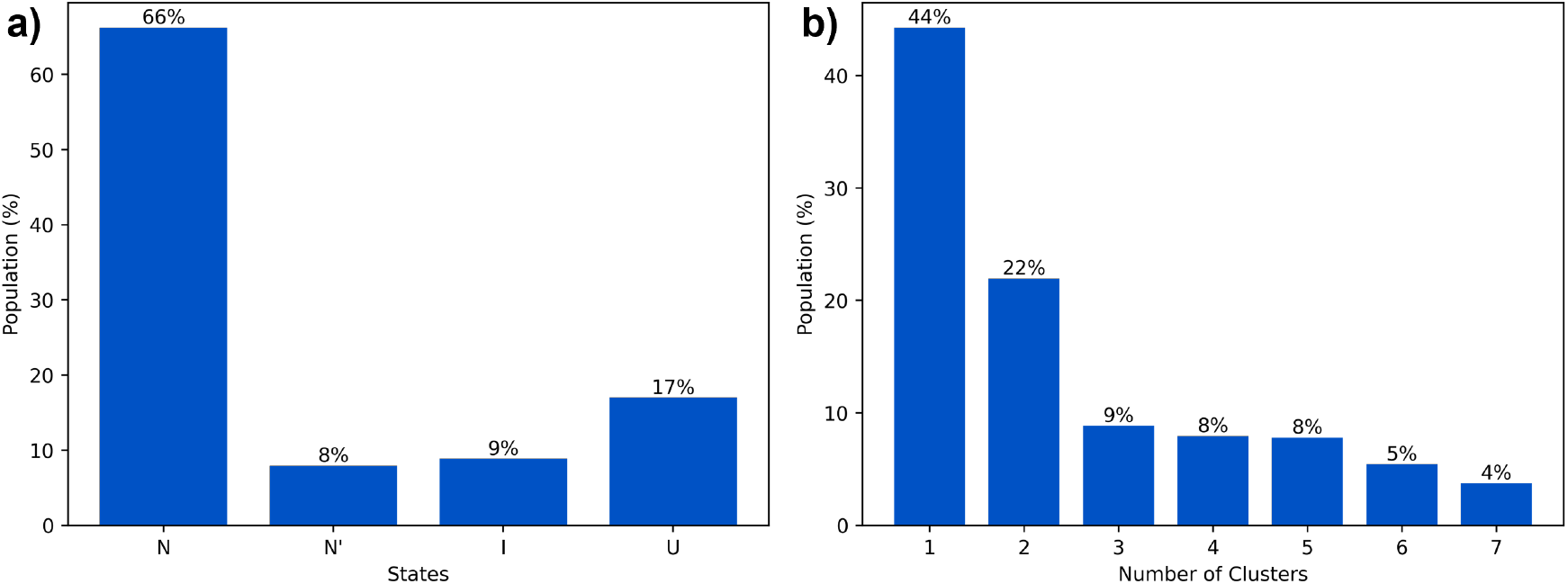
**(a)** Cluster population for HP35 in four states: folded (N), partially folded (N’), inter-mediate (I), and unfolded (U). **(b)** Individual cluster population.

**Figure 11:**
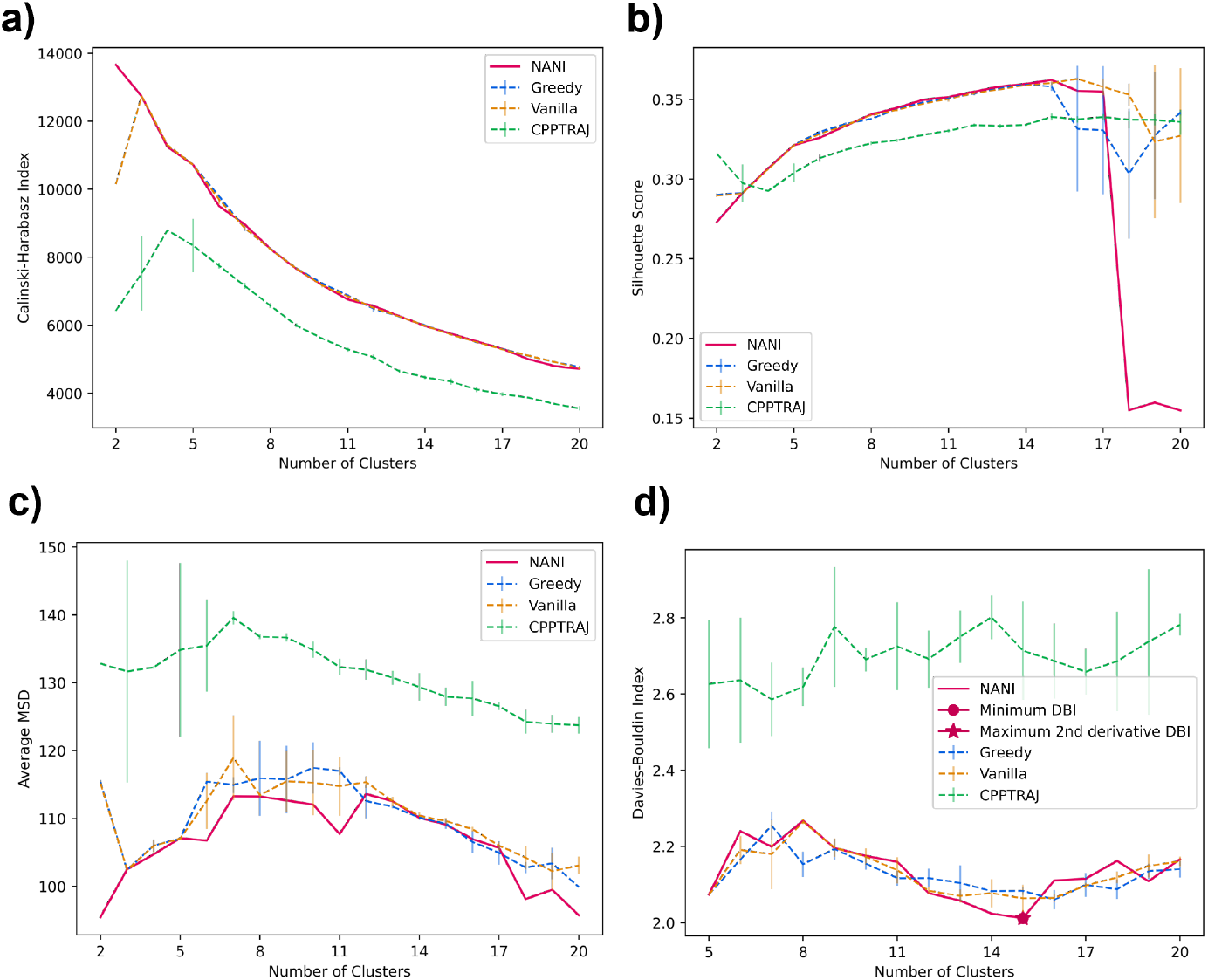
Indicators of different seed selectors applied on NuG2. Error bars represent the standard deviation over three replicates. All use the single-reference alignment. **(a)** Calinski-Harabasz index **(b)** Silhouette score **(c)** Average mean squared deviation **(d)** Davies–Bouldin index

**Figure 12:**
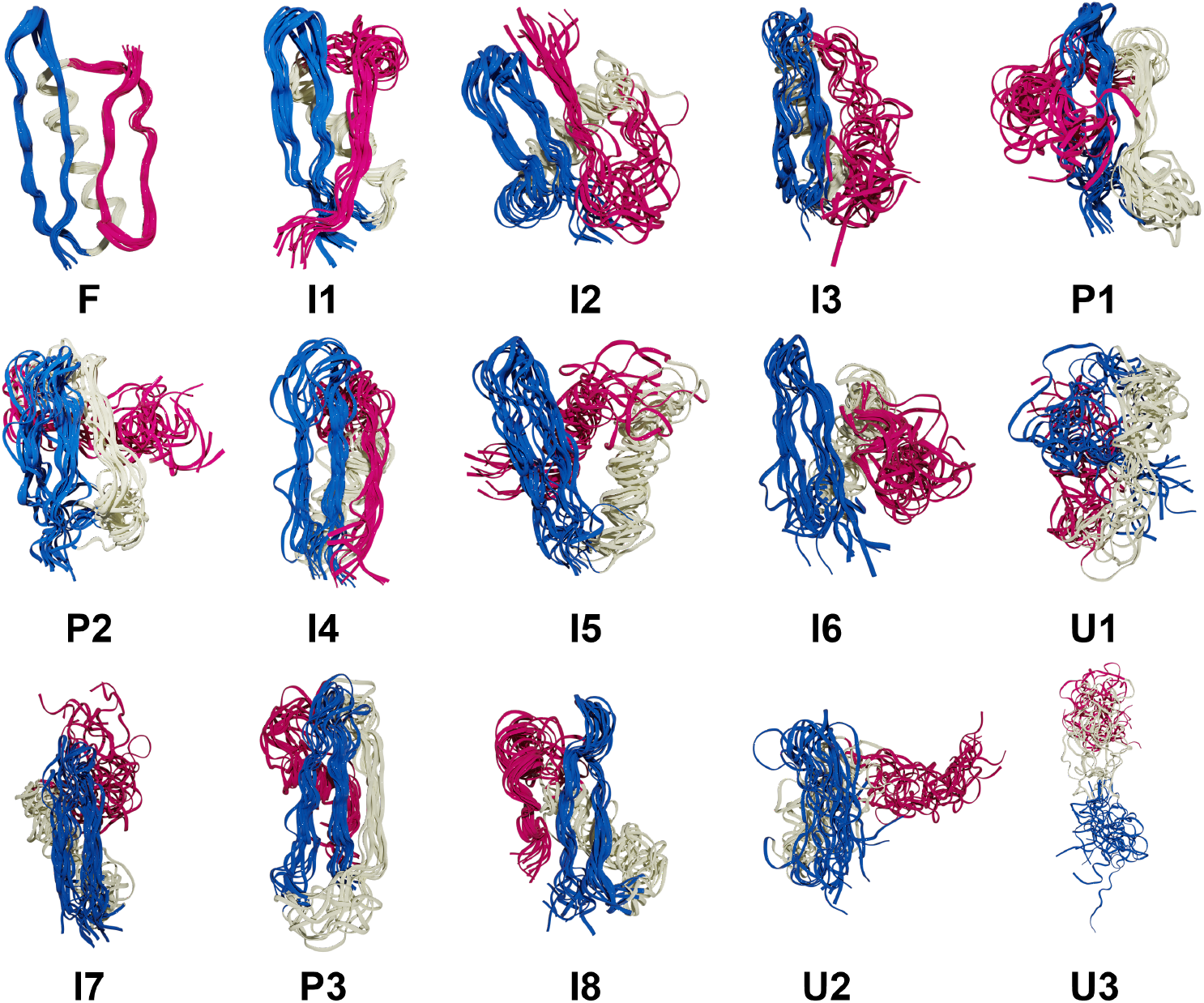
Structural overlaps for NuG2 in four states: folded (F), intermediate (I), pre-folded (P), and unfolded (U). *β*1 and *β*2 strands are colored blue, the helix is colored white, and *β*3 and *β*4 strands are colored pink.

**Figure 13:**
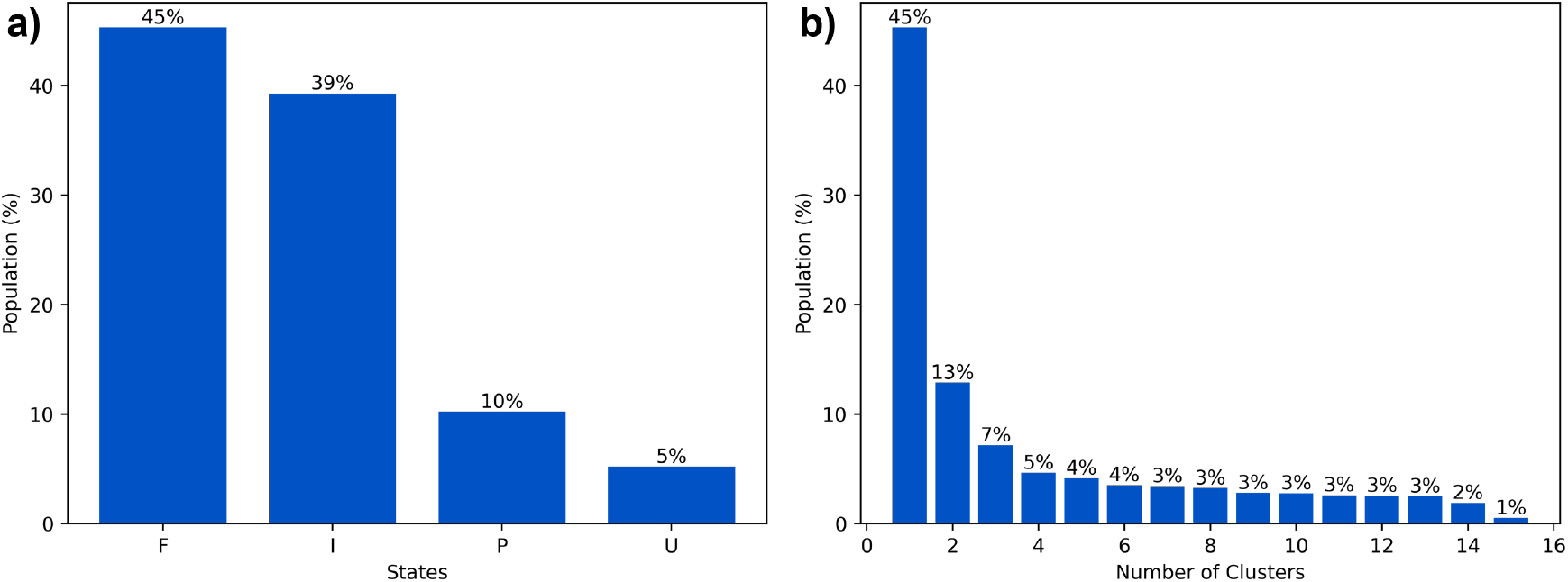
**(a)** Cluster population for NuG2 in four states: folded (F), intermediate (I), pre-folded (P), and unfolded (U). **(b)** Individual cluster population.

Kronecker alignment has some limitations compared to single-reference alignment (See Fig. S7) because it is unable to recognize the N’ state. This is surprising since this alignment was able to find this conformational basin when combined with Shape-GMM clustering. This seems to indicate that for a simple method, like *k*-means NANI, the single reference alignment might be a more convenient starting point.

### NuG2

NuG2 has a complex protein-folding mechanism as it has two *β*-sheets and an *α*-helix, and can follow two different pathways to fold.^60^ Given the results of the previous section, the clustering for NuG2 only uses the single-reference alignment. The same trend from the peptide section is also observed in NuG2 clustering: NANI can generate compact and well-defined clusters as shown in the lower MSD and DBI values. The results from all flavors of the *k*-means++ predecessors fluctuated greatly between replicates, significantly altering the number of clusters per run. Once again, the standard CPPTRAJ *k*-means implementation resulted in the highest MSD and DBI values and the lowest CH values, while also showing a significant variance in the final results, like the *k*-means++ flavors. The number of clusters was determined to be fifteen calculated using NANI as the seed selector. The folded structure was the predominant cluster at 45%, eight intermediate structures were identified at 39%, three pre-folded structures were found at 10%, and lastly, three unfolded structures were observed at 5%. It is remarkable that an algorithm as simple and efficient as NANI was able to find metastable states corresponding to both folding pathways. In our previous studies on this system, we showed that traditional hierarchical agglomerative clustering (HAC) methods failed to identify several intermediate conformations along the folding process.^20^ Only a variant of HAC that used an *n*-ary linkage criterion (and was based on a contact map representation of NuG2) was able to distinguish between these states. However, as with every HAC algorithm, this (at best) scales as *O*(*N*^2^), so it is markedly less efficient than *k*-means NANI. The dominant folded structure revealed well-overlapped structures in the cluster, with the lowest variances. In all the intermediate, a partial unfolding of the *β*3/4 strands is observed; due to this unfolding, many different conformations were adapted by *β*3/4. In all the pre-folded states, all *β*3/4 and the *α*-helix were partially unfolded; moreover, *β*1/2 was also observed to have greater variance in the structure compared to the structures in the intermediate states. Unfolded states uncovered a complete unfolding of all three secondary structures in the protein.

## Conclusion

NANI is a robust seed selector for *k*-means that surpasses most publicly available seed selectors such as *k*-means++ and the seed selector for CPPTRAJ *k*-means. Its advantage comes from its reproducibility as a deterministic algorithm, which gives the same optimal number of clusters and cluster population at each run. Furthermore, as shown in the lower DBI and average MSD values, it excels at creating tight and well-defined clusters. When applied to model 2D systems, the `comp sim` version of NANI (the one that chooses centroids only from previously stratified data) highlighted the cause of the potential issues found in the traditional *k*-means++ and CPPTRAJ implementations. That is, the diversity-picking algorithms applied to the whole set of points tend to select initial centroids in the boundary of the set, which are typically low-density regions. Hence, even though it is expected that the final centroids are well-separated, they will also be found in regions with a higher density of frames. This is more evident with the standard CPPTRAJ implementation, which is the one that most aggressively tries to maximize the separation between initial centroids, which seems to cause the optimization of the *k*-means objective function to be stuck into local minima where the final centroids are not optimally distributed. Both *k*-means++ initializations tend to correct this behavior by using a probabilistic diversity exploration mechanism. In this way, they de-emphasize the role of very distant points and allow themselves to potentially select guesses in the high-density zones. However, this introduces further randomness in the final results. NANI combines the best of these worlds, by trying to maximize the diversity of the initial centroids but limiting the exploration of a pre-stratified subset of the data, that is more likely to contain only regions with a higher density of points. The study of even simple peptide systems showcased one of the key issues of any clustering study: identifying the optimum number of clusters in the data. We considered several popular clustering evaluation metrics for doing so, including the CH index and Silhouette score. The former showed an almost perfect monotonically increasing tendency with decreasing number of clusters, while the latter was either virtually constant over ranges of *k* values, or also monotonically increasing (or, in the case of the *β*-hairpin, its results varied wildly depending on the alignment). On the other hand, the DBI clustering evaluation metric proved to be more amenable to interpretation. While it can also show a marked bias towards very small *k* values, we found two ways to work around this issue. First, the overall non-monotonic behavior of this index means that if one restricts the scan of *k* values to not include very small *k* values (*k* = 2, 3, 4), it is possible to identify a physically meaningful number of clusters. Moreover, if one decides to follow the local behavior of the DBI index, instead of its absolute value, and uses the 2^nd^ derivative as a way to gauge the relative stability of the *k* values, it is also possible to determine the optimum *k* values. Finally, on the matter of preferred *k* values, while the absolute values of the average MSD cannot be used for this purpose, we noted that the 2^nd^ derivative of the MSD values correlates with the ideal *k* values found using the DBI index. Given the ease of calculation of the MSD compared to the DBI, it seems reasonable to perform an MSD scan analysis over extended ranges of *k* ‘s, and then only perform the DBI calculations for the *k* values identified through the MSD analysis. This seems like a convenient strategy to aid in speeding up the post-processing of the clustering results, which we will explore in a forthcoming contribution.

The MD simulations also served to highlight some of the key advantages of NANI. Above all, NANI offers a reproducibility that is not attainable with the other *k*-means algorithms. While the probabilistic nature of the *k*-means++ methods does not have a great impact on the final results for model 2D systems, this quickly changes even for simple MD simulations. The CH, MSD, and Silhouette scores profiles for both *k*-means++ flavors showed marked differences from one run to the other, and this was particularly evident for the DBI. This is especially concerning since, as discussed before, DBI seems to be the more robust at the time of determining the ideal *k* for the given simulation, i.e. different *k*-means++ runs can lead to different answers about the number of clusters present in the data. The CPPTRAJ *k*-means also shows great variability, but it is also accompanied by overall lower quality indicators of the considered scores. This indicates that even if we were to use some deterministic criterion to choose the initial centroid in the CPPTRAJ algorithm, choosing the remaining *k* − 1 initial centers doing a diversity screening over all the frames will lead to picking centers in very low-populated regions in the data, which is a sub-optimal choice. (This agrees with the comparatively sub-par performance of the `div select` algorithm on the model 2D systems.) NANI then offers a fully deterministic alternative, with overall performance on par (if not better) than the *k*-means++ alternatives. We also studied two different alignment paradigms: the traditional single-reference, and the Hocky and McCullagh Kronecker alignments.^35^ We observed that single-reference tends to produce more compact clusters (as measured by the average MSD), while the Kronecker method tends to result in better-separated clusters (as quantified by the DBI). In general, both alignment methods have strengths and weaknesses, and it is certainly desirable to explore their relative performance when compared in conjunction with other clustering algorithms. In the present case, we tend to slightly favor the combination of single-reference with *k*-means because this was the combination that allowed identifying the previously reported metastable states for the HP35 and NuG2 systems. It is reassuring to see how NANI was able to not only find the previously identified N, N’, I, and U states of HP35 but also with relative populations in good agreement with those found by more complex (and time-consuming) algorithms. The same success was found for the NuG2 protein, with NANI being able to identify several metastable states across multiple folding pathways that were absent even in the (markedly) less efficient hierarchical studies.

This study introduced NANI as the first installment of our MDANCE clustering package, which can be used as a standalone software or part of our clustering package. Aside from introducing novel clustering algorithms, a pipeline for clustering analysis is also available for determining the most optimal number of clusters for the dataset. In forthcoming contributions, we will expand on the functionality of MDANCE, and we will add more clustering algorithms to this package based on the use of *n*-ary similarity and difference functions.

## Acknowledgement

We thank Martin McCullagh for his help with the ShapeGMM package. RAMQ and LC thank support from the National Institute of General Medical Sciences of the National Institutes of Health under award number R35GM150620. CS and MK thank National Institutes of Health for support through grant GM107104.

## Notes

https://github.com/mqcomplab/MDANCE

### Competing Interest Statement

The authors have declared no competing interest.

