## Supplementary Information for "k-Means NANI: an improved clustering algorithm for Molecular Dynamics simulations"

### Supporting Information Available

#### 2D Datasets

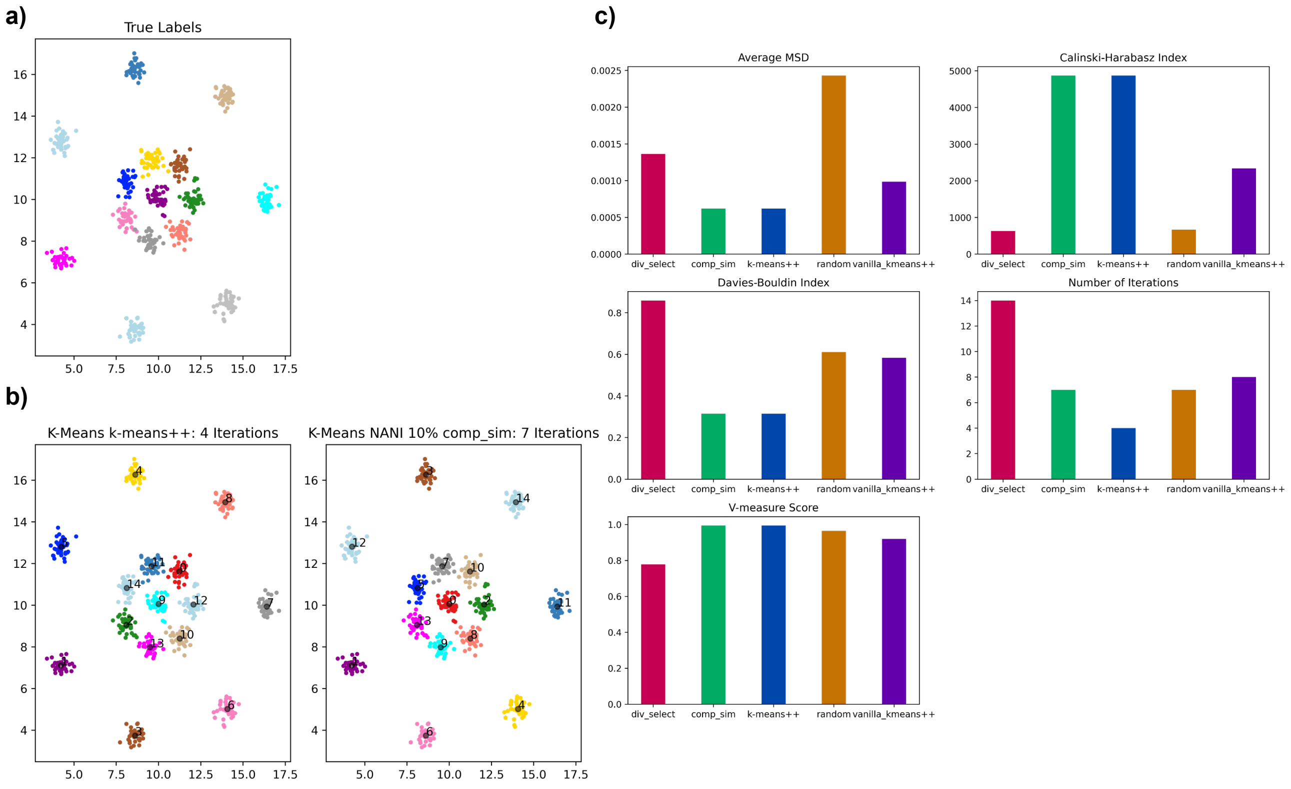


Figure S1: $k$-means NANI on a sample blob disk data. A different color represents a different cluster label. The black dot represents the centroid of that cluster. **(a)** True labels. **(b)** $k$-means clustering with centroids initialized by $k$-means++ versus NANI. **(c)** Summary of average MSD, Calinski-Harabasz index, Davies-Bouldin index, number of iterations, and V-measure score using different seed selectors.

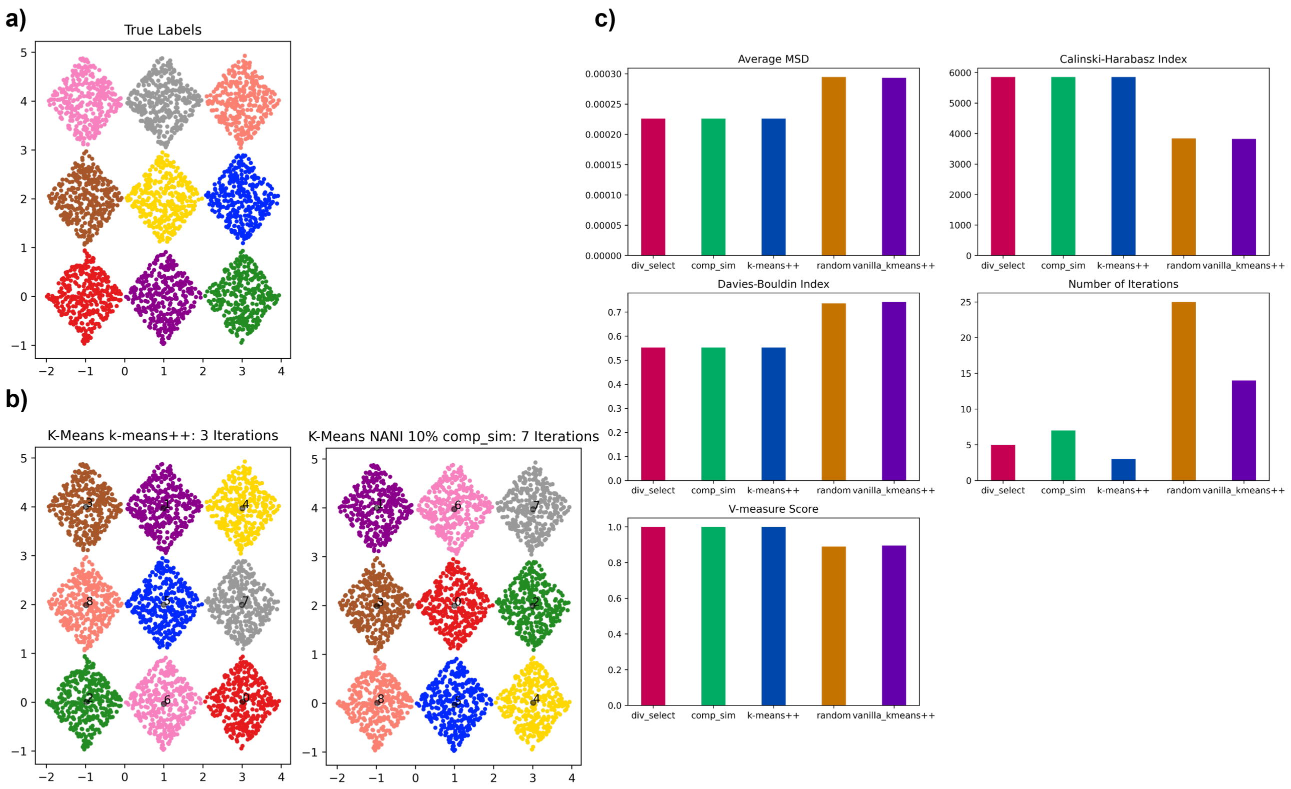


Figure S2: $k$-means NANI on a sample diamonds data. A different color represents a different cluster label. The black dot represents the centroid of that cluster. **(a)** True labels. **(b)** $k$-means clustering with centroids initialized by $k$-means++ versus NANI. **(c)** Summary of average MSD, Calinski-Harabasz index, Davies-Bouldin index, number of iterations, and V-measure score using different seed selectors.

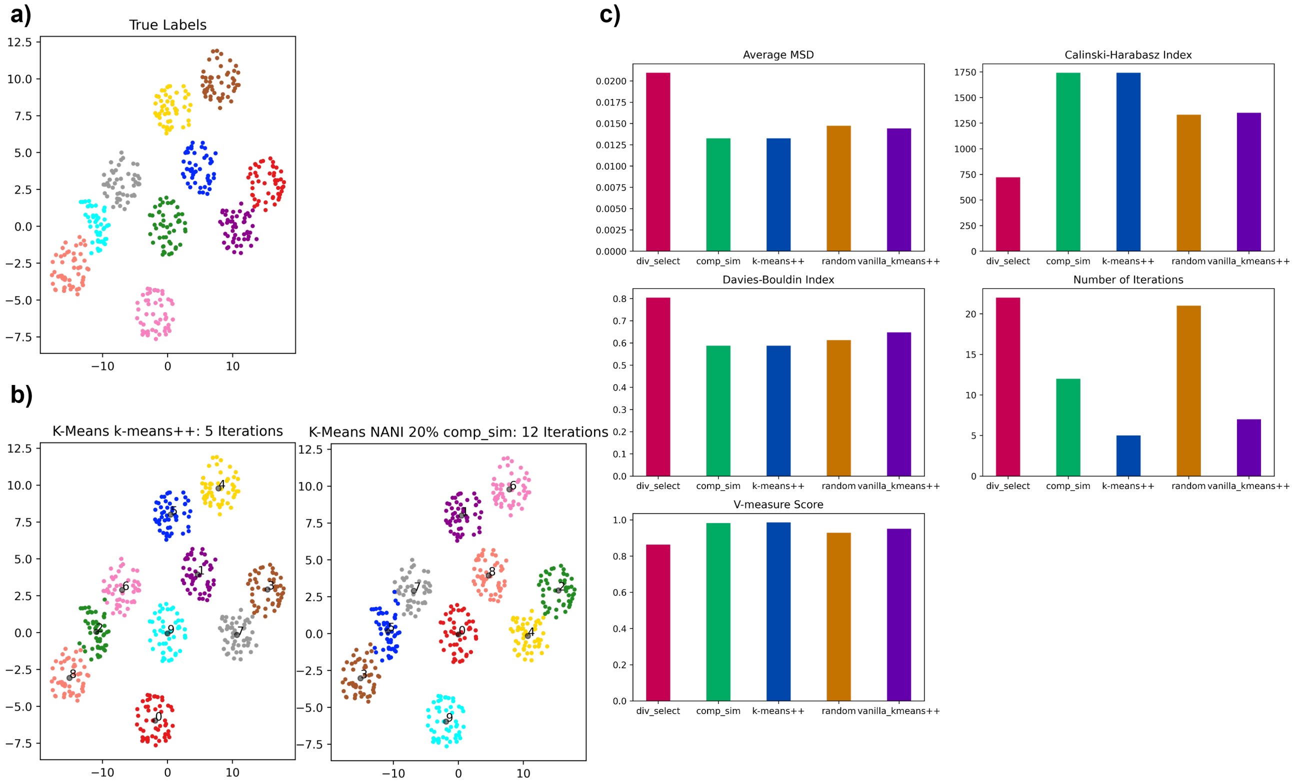


Figure S3: $k$-means NANI on a sample ellipses data. A different color represents a different cluster label. The black dot represents the centroid of that cluster. **(a)** True labels. **(b)** $k$-means clustering with centroids initialized by $k$-means++ versus NANI. **(c)** Summary of average MSD, Calinski-Harabasz index, Davies-Bouldin index, number of iterations, and V-measure score using different seed selectors.

### Peptides

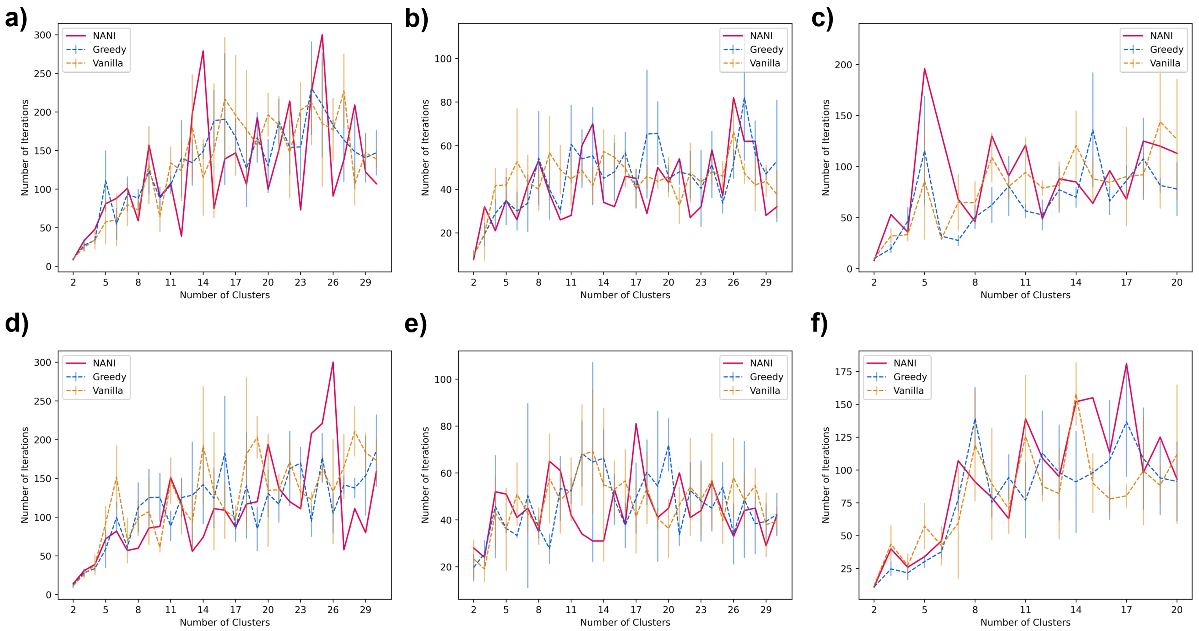


Figure S4: Number of Iterations of different seed selectors applied on three peptide systems. **(a)** R1Q hairpin single-reference aligned **(b)** $\beta$-heptapeptide single-reference aligned **(c)** Villin single-reference aligned **(d)** R1Q hairpin Kronecker aligned **(e)** $\beta$-heptapeptide Kronecker aligned **(f)** Villin Kronecker aligned

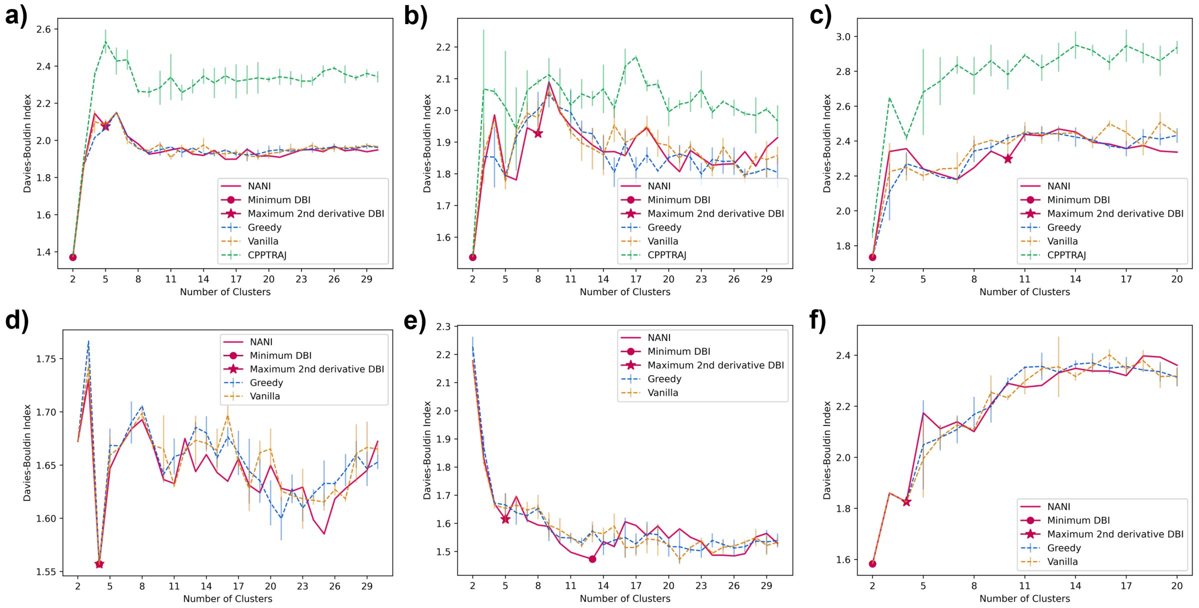


Figure S5: Davies-Bouldin Index of different seed selectors applied on three peptide systems starting at 2 clusters. **(a)** R1Q hairpin single-reference aligned **(b)** $\beta$-heptapeptide single-reference aligned **(c)** Villin single-reference aligned **(d)** R1Q hairpin Kronecker aligned **(e)** $\beta$-heptapeptide Kronecker aligned **(f)** Villin Kronecker aligned

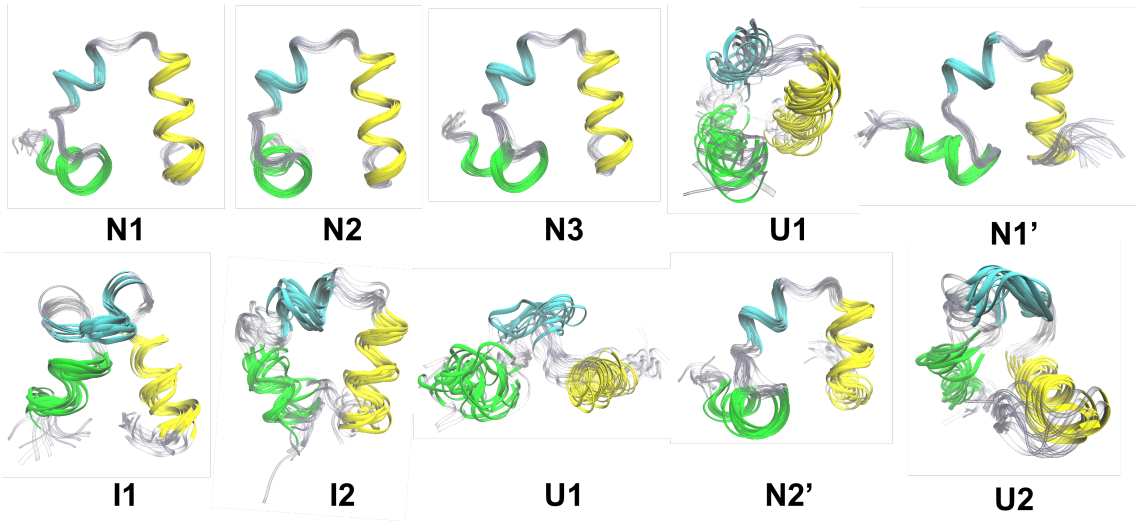


Figure S6: Structural overlaps for Villin in four states: folded (N), partially folded (N'), intermediate (I), and unfolded (U). Helix 1, 2, and 3 are in green, cyan, and yellow, respectively. Clustering was done using single-reference alignment and ten clusters were the maximum $2^{nd}$ derivative DB determined from DB plots in Fig. 7c.

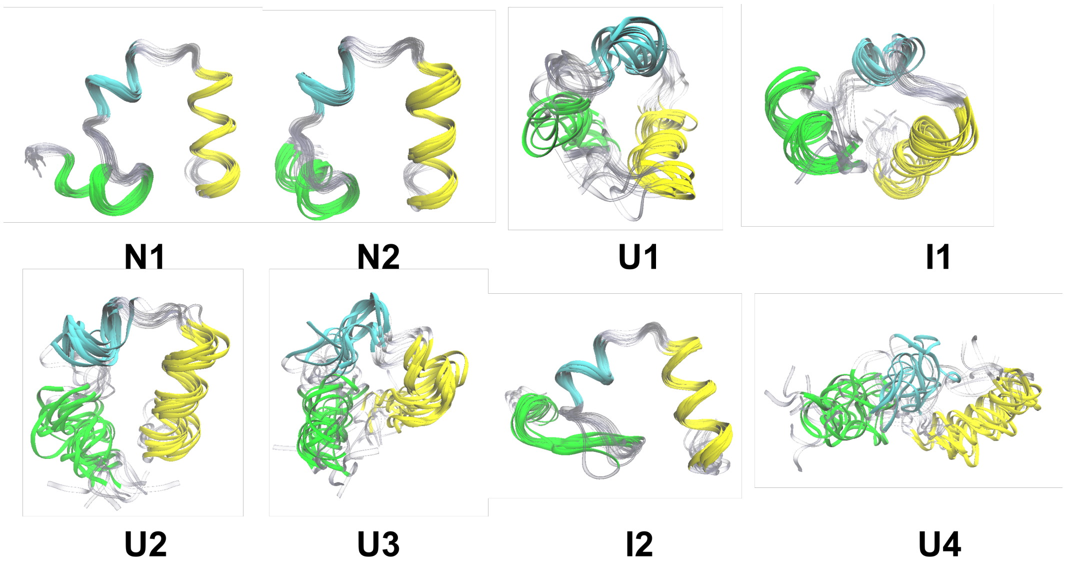


Figure S7: Structural overlaps for Villin in three states: folded (N), intermediate (I), and unfolded (U). Helix 1, 2, and 3 are in green, cyan, and yellow, respectively. Clustering was done using Kronecker alignment and eight clusters were the minimum DB determined from DB plots in Fig. 7f.

#### Protein

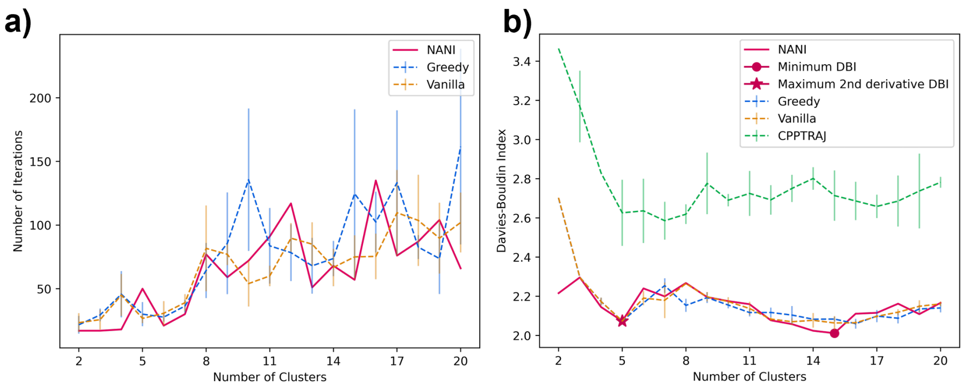


Figure S8: Indicators of different seed selectors applied on NuG2. **(a)** Number of iterations **(b)** Davies-Bouldin index starting at 2 clusters
